# FACT depletion results in a temporal cascade of chromatin disruption preceding transcriptional collapse in stem cells

**DOI:** 10.64898/2026.02.26.708215

**Authors:** Rithika Sankar, Bryona M. Jackson, Santana M. Lardo, Valentin Flury, Anja Groth, Sarah J. Hainer

## Abstract

The histone chaperone FACT is essential for chromatin organization and stem cell identity, yet the sequence of events through which FACT maintains chromatin integrity remains unclear. We generate a high-resolution temporal map of events leading to chromatin destabilization and cell identity loss in murine embryonic stem cells following dTAG-mediated depletion of FACT. Loss of FACT initiates a rapid cascade, with nucleosome phasing disruption and reduced H3K4me3 beginning within 10 minutes in a transcription-dependent manner. This architectural destabilization enables pluripotency transcription factors to ectopically occupy gene bodies. FACT loss results in RNA Polymerase II accumulation within 30 minutes and altered H3K36me3 deposition preceding a global transcriptional collapse at 2 hours of FACT depletion. FACT restoration fails to rescue these defects, demonstrating they are largely irreversible. Our findings define a temporal hierarchy in which FACT loss drives progressive deterioration of chromatin architecture, leading to transcriptional collapse in stem cells.

## INTRODUCTION

The packing of DNA into chromatin serves not only to compact genetic material within the nucleus, but also to regulate access to the underlying sequence. Chromatin is dynamically remodeled to permit or restrict the activity of DNA-templated processes, including transcription^1,2^. Consequently, precise regulation of chromatin organization is essential for appropriate gene expression and cell fate decisions^3^. Nucleosomes, octameric complexes of histone proteins around which ∼147 bp of DNA is wrapped, present a significant barrier to transcription by RNA Polymerase II (RNAPII)^4,5^. For RNAPII to access and traverse the DNA template, nucleosomes must be transiently displaced and subsequently reassembled to maintain chromatin integrity^6–8^. This dynamic cycle of nucleosome disassembly and reassembly is orchestrated by nucleosome remodelers and histone chaperones that act co-transcriptionally to facilitate RNAPII passage while preserving epigenetic information^9–11^. Among these factors, the FACT (FAcililates Chromatin Transcription) complex serves as a central regulator of transcription-coupled nucleosome dynamics^12–14^.

FACT is a highly conserved and essential histone chaperone, first identified as a factor required for transcription elongation through chromatin templates *in vitro*^15–17^. Recent *in vitro* studies have greatly advanced our understanding of FACT activity. First, FACT is only able to act on partially disrupted nucleosomes, and these nucleosomes are likely unraveled by the transcription machinery and other factors such as nucleosome remodelers. FACT is able to then displace histones and simultaneously transfer dimers upstream of RNAPII, promoting reassembling of nucleosomes,^18–22^. Together these data support an important role for FACT in transcription elongation and maintenance of the chromatin landscape.

In *S. cerevisiae*, FACT predominantly occupies actively transcribed genes to facilitate RNAPII passage ^20,22–27^. There, FACT is required for recycling of histones which is critical for preserving the chromatin landscape over actively transcribed genes. Consistently, studies in *S. cerevisiae* and *D. melanogaster* have demonstrated that FACT is required to maintain histone modifications over transcribed regions, including the promoter-proximal mark H3K4me3 and the transcription elongation-associated mark H3K36me3^28–31^. Functional studies in *D. melanogaster* have also implicated FACT in the regulation of RNAPII promoter-proximal pausing, highlighting its influence beyond transcription elongation^31^.

Beyond its canonical role in transcription elongation, FACT has been implicated in the maintenance of cellular identity in mammals^32–39^. The mammalian FACT complex is a heterodimer composed of the dimer exchange subunit SPT16 (Suppressor of Ty 16 homolog) and an HMG-containing subunit SSRP1 (Structure-Specific Recognition Protein 1)^16^. In mammalian cells, FACT occupies gene bodies of actively transcribed genes as well as promoters and enhancers^31,32,38,39^. FACT is highly expressed in undifferentiated cell types, including stem cells and cancer cells, where it is essential for maintaining cellular identity and proliferative capacity^34,36,39^. In murine embryonic stem (ES) cells, FACT depletion leads to loss of pluripotency, accompanied by reduced expression of pluripotency transcription factors (TFs), including *Oct4*, *Sox2*, and *Nanog*, and altered nucleosome occupancy and chromatin accessibility^38^. Similarly, numerous cancer cell types depend on FACT for survival, and FACT inhibition is actively being explored as a therapeutic strategy^40–44^. Although FACT’s contributions to chromatin regulation and cell identity have been studied, the precise mechanism by which FACT-dependent chromatin organization maintains cell fate remains unclear.

To address the direct mechanism of FACT in preserving cell identity, we developed two rapid, inducible degron TAG (dTAG)-mediated depletion systems targeting either SPT16 or SSRP1 in murine ES cells. This approach enables precise temporal resolution of molecular events linking FACT loss to chromatin destabilization and cell fate changes. FACT depletion causes rapid disruption of nucleosome phasing and reduction of H3K4me3 at the 5’ ends of actively transcribed genes, observed through CUT&RUN and MNase-seq and detectable as early as 10 minutes post-depletion. Next, RNAPII accumulates at the 5’ ends of genes by 30 minutes post-depletion and we then observe a reduction in H3K36me3, indicative of an elongation defect. In parallel, pluripotency transcription factors (OCT4, SOX2, NANOG) invade into gene bodies. This ectopic occupancy correlates with transcriptional output and occurs at sites of chromatin disruption and independent of motif preferences, suggesting that FACT normally maintains the physical barriers that restrict TF access. Single-molecule Fiber-seq and FIRE analysis corroborate nucleosome loss observed by short-read approaches and suggest that TFs and RNAPII accumulation occur on the same DNA molecules. These chromatin changes ultimately culminate in widespread transcriptional dysregulation, which are not observed until 2 hours post depletion, establishing that chromatin disruption drives, rather than accompanies, transcriptional collapse. Furthermore, transient FACT depletion causes epigenomic alterations that cannot be fully reversed upon FACT restoration, underscoring the fragility of the chromatin landscape and demonstrating that FACT-dependent chromatin organization is essential for maintaining ES cell identity.

## RESULTS

### Rapid depletion of FACT results in increased transcription factor occupancy over gene bodies

To examine the precise mechanism through which FACT regulates pluripotency in ES cells, we acutely depleted the FACT subunits SPT16 and SSRP1 using the dTAG system in murine ES cells grown in naïve conditions (serum/LIF + 2i; Fig. 1A). Relative to vehicle control (DMSO), addition of dTAG-13 (referred throughout as dTAG) resulted in rapid depletion of SPT16 protein starting at 10 minutes of treatment (Fig. 1B-C, S1A-D). Both SPT16 and SSRP1 were fully depleted within 2 hours of dTAG treatment. As observed previously, SPT16 or SSRP1 depletion led to a co-depletion of the other complex subunit (Fig. S1A-D)^45^. Consistent with a role for FACT in stem cell identity and viability, we observed a reduction in cell viability starting at 6 hours of SPT16 depletion and 12 hours of SSRP1 depletion (Fig. S1E).

**Figure 1.**
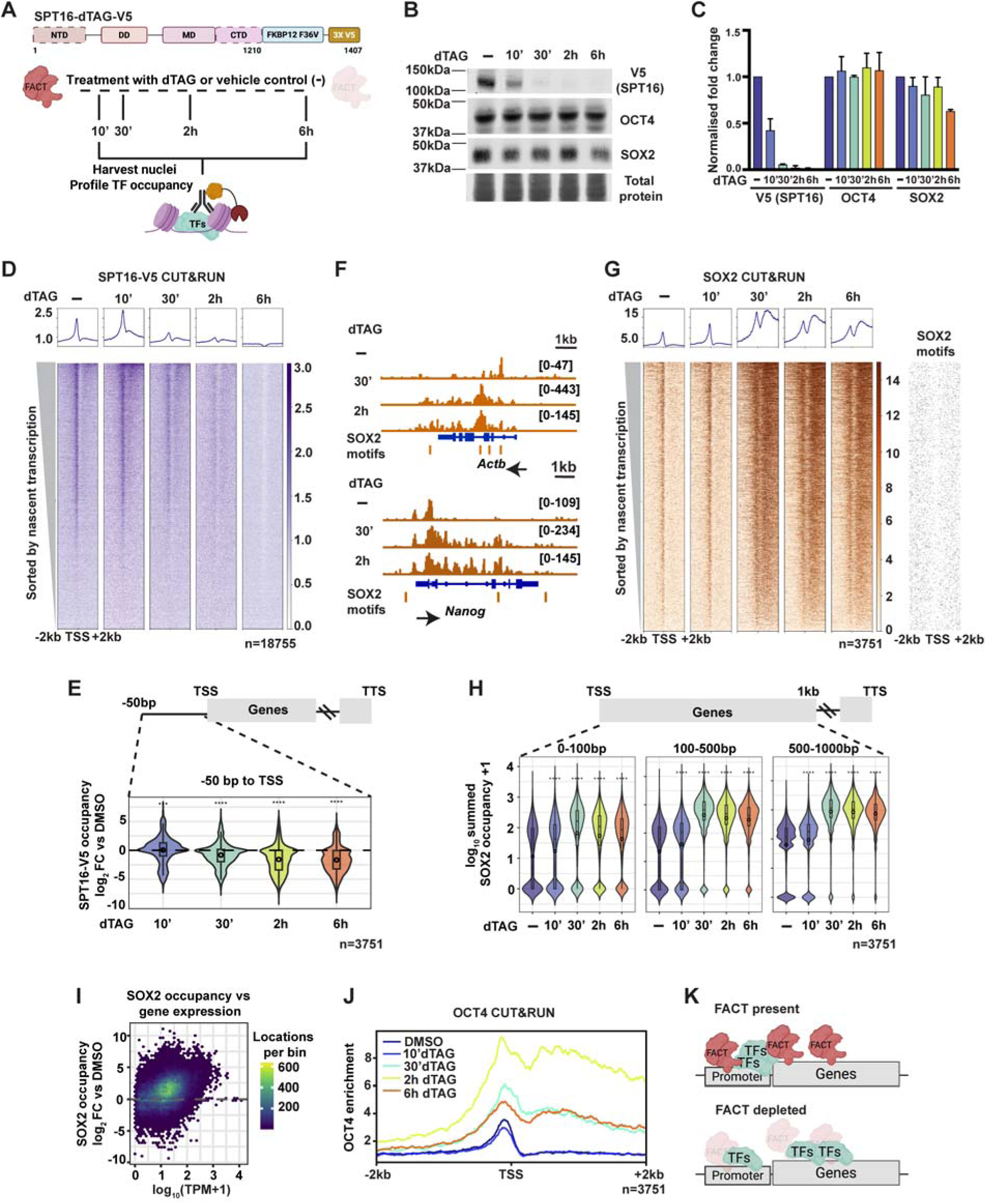
Acute FACT depletion causes transcription factor mislocalization into gene bodies. **(A)** Schematic of the SPT16-dTAG-V5 system (referred to as SPT16-dTAG throughout). The endogenous *Supt16h* locus was modified to include FKBP12 (F36V) and 3X-V5. Murine ES cells were treated with dTAG-13 or vehicle control (−; DMSO) for indicated times, followed by nuclei isolation and CUT&RUN to profile transcription factor (TF) occupancy. **(B)** Western blot analysis of SPT16-V5, OCT4, and SOX2 protein levels following dTAG-13 treatment in SPT16-dTAG ES cells. Total protein stained with REVERT serves as a loading control. **(C)** Quantification of protein levels, normalized to total protein. Data represent normalized fold change ± SD. **(D)** Heatmap of SPT16-V5 CUT&RUN signal over transcription start sites (TSSs ± 2 kb) performed in SPT16-dTAG ES cells across the dTAG-13 treatment time course. Genes (n = 18,755) are sorted by nascent transcription levels from untreated cells^83^. Metaplot (top) shows average signal across all genes. **(E)** Quantification of SPT16-V5 occupancy changes (log_₂_ fold change versus DMSO) within 50 bp upstream of the TSS for highly transcribed genes (n = 3,751). Top quintile of genes were selected as highly transcribed throughout the manuscript. ****p < 0.0001 (For each genomic window summed occupancy was compared across conditions using a Kruskal–Wallis test. Pairwise differences relative to DMSO were assessed using two-sided Wilcoxon rank-sum tests with Bonferroni correction; significance is indicated by asterisks). **(F)** Averaged genome browser tracks showing SOX2 CUT&RUN signal and SOX2 motifs over the *Actb* and *Nanog* loci across treatment conditions in SPT16-dTAG ES cells. Orange bars below gene indicate SOX2 motifs. **(G)** Heatmap of SOX2 CUT&RUN signal at TSSs ± 2 kb across the dTAG-13 treatment time course in SPT16-dTAG ES cells. Top quintile of transcribed genes (n = 3,751) is shown and sorted by nascent transcription levels^83^. Metaplot (top) shows average signal. Right: Heatmap showing positions (binary, black signal indicates presence) of SOX2 motifs over TSSs ± 2 kb of top quintile genes (n=3,751). **(H)** Violin plots showing summed SOX2 occupancy (log_₁₀_+ 1) across three gene body regions (0-100 bp, 100-500 bp, and 500-1000 bp from TSS) across treatment conditions in SPT16-dTAG ES cells. ****p < 0.0001 (For each genomic window summed occupancy was compared across conditions using a Kruskal–Wallis test. Pairwise differences relative to DMSO were assessed using two-sided Wilcoxon rank-sum tests with Bonferroni correction; significance is indicated by asterisks). **(I)** Density scatterplot showing the relationship between SOX2 occupancy changes (log_₂_ fold change versus DMSO) and gene expression levels (log_₁₀_TPM + 1) at 30 min SPT16-dTAG depletion. Color scale indicates the number of genomic locations per bin. **(J)** Metaplot of OCT4 CUT&RUN signal at TSSs ± 2 kb across treatment conditions in SPT16-dTAG ES cells for highly transcribed genes (n = 3,751). **(K)** Model depicting FACT impact on TF localization. When FACT is present (top), TFs are appropriately localized to promoters. Upon FACT depletion (bottom), TFs redistribute into gene bodies. See also Figure S1 and Figure S2.

We examined chromatin localization of FACT using Cleavage Under Target and Release Using Nuclease (CUT&RUN) against the V5 epitope appended to SPT16 and observed a progressive reduction in SPT16 chromatin occupancy over the depletion time course (Fig. 1D). Quantification of SPT16-V5 occupancy within 50 bp upstream of the transcription start site (TSS) revealed that among the top quintile of transcribed genes, ∼40% showed reduced SPT16-V5 occupancy relative to DMSO as early as 10 minutes post-depletion (Fig. 1E). We also visualized FACT occupancy over previously established FACT binding sites and confirmed a reduction in occupancy throughout the depletion time course (Fig. S1F)^38^.

We previously reported that FACT depletion led to a progressive reduction of pluripotency factor expression, resulting in rapid differentiation and cell death in ES cells^38^. Since those experiments were performed with a slow-depleting SPT16-AID ES cell line, we asked whether pluripotency gene expression was similarly downregulated upon rapid dTAG-mediated depletion. Indeed, mRNA levels of pluripotency factors were downregulated within 2 hours of SSRP1 depletion (Fig. S1G). Moreover, ∼60% of SPT16 peaks overlap with OCT4, SOX2, and/or NANOG binding sites^38^. Based on these observations, we hypothesized that FACT might be directly responsible for appropriate localization of pluripotency-relevant TFs at their target sites. Because OCT4, SOX2, and NANOG positively regulate their own expression, we predicted that disrupted TF localization upon FACT depletion would lead to downregulation of pluripotency gene expression^46,47^. To test this, we depleted FACT and performed CUT&RUN to assess TF occupancy. Contrary to our prediction, FACT depletion did not reduce TF occupancy at target sites. Rather, OCT4, SOX2, and NANOG accumulated within gene bodies upon FACT depletion (Fig. 1F-H, Fig. S1H-J, S2A). We observed an increase in SOX2 occupancy over gene bodies starting at 30 minutes of SPT16 depletion and by 2 hours of SSRP1 depletion (Fig. 1G-H). To orthogonally assess TF localization upon FACT depletion, we performed MNase-ChIP to measure genome-wide OCT4 and SOX2 occupancy. Relative to vehicle control-treated samples, we observed increased gene body occupancy of OCT4 and SOX2 in both SPT16 and SSRP1 2 hour depletion conditions (Fig. S2B). Notably, SPT16 depletion resulted in TF mislocalization as early as 30 minutes post-treatment, whereas SSRP1 depletion resulted in similar changes by 2 hours post-treatment. This temporal difference may reflect the distinct roles of each subunit in FACT complex function and stability.

Since FACT occupancy correlates with transcription (Fig. 1D), we predicted that highly transcribed genes would exhibit the most pronounced mislocalization of TF occupancy. Consistent with this prediction, when SOX2 occupancy is sorted by nascent transcription, highly transcribed genes showed the greatest increase in TF occupancy over the gene body (Fig. 1G). We quantified summed SOX2 occupancy over different bins within 1 kb of the TSS and observed an increase in occupancy over the time course of SPT16 depletion (Fig. 1H). Similarly, plotting SOX2 gene body occupancy as a function of gene expression revealed a positive correlation between SOX2 occupancy and transcriptional output (Fig. 1I).

Given that SOX2 protein levels were not reduced until 6 hours of FACT depletion (Fig. 1B-C, S1A-B), we asked whether TFs were being redistributed away from their canonical genomic binding sites. To address this, we called peaks from SOX2 CUT&RUN data in control and depleted conditions. We found an increase in the proportion of genic peaks, along with a progressive reduction in the proportion of peaks over intergenic, promoter, and repetitive elements through the depletion time course (Fig. S2A). These data suggest that SOX2 is potentially redistributed from canonical binding sites to gene bodies. To assess whether SOX2 occupancy was moving into genic locations based on motif preferences, we visualized SOX2 motifs ±2 kb from the TSS over highly transcribed locations. We observed a relatively even distribution of SOX2 motifs over the 4 kb window, with no correlation between motif density and ectopic SOX2 occupancy downstream of the TSS (Fig. 1G). We also observed no enrichment of motifs specifically over the highly transcribed genes, where mislocalized SOX2 occupancy is greatest. Furthermore, since we also observed mislocalization of NANOG (Fig. S1J), we conclude that this phenomenon is not restricted to TFs previously described as pioneer factors, but rather likely affects many DNA-binding proteins independent of motifs. Together, these results demonstrate that FACT depletion causes redistribution of TFs into gene bodies, suggesting that FACT normally functions to restrict TF occupancy to appropriate regulatory elements (Fig. 1K).

### FACT depletion leads to reduced nucleosome phasing and H3K4me3 occupancy over the 5’ ends of genes, preceding TF mislocalization

Having established that FACT depletion results in TF mislocalization into gene bodies at 30 minutes (SPT16-dTAG) or by 2 hours (SSRP1-dTAG) of depletion, we next asked whether disruption of chromatin architecture might permit this promiscuous TF occupancy. To assess this, we depleted FACT and profiled the histone modification H3K4me3 using CUT&RUN, through which we can also infer nucleosome positions, and assessed nucleosome occupancy using MNase-seq (Fig. 2A). Examination of individual loci, such as the *Nanog* gene, revealed a progressive loss of H3K4me3 signal beginning by 10 minutes of SPT16 depletion (Fig. 2B). Notably, total H3K4me3 protein levels were unchanged over the depletion time course (Fig. S2C-D). More globally, by 10 minutes of SPT16 depletion, H3K4me3 reduction was observed over the 5’ ends of highly transcribed genes (Fig. 2C-E, S2E-F, S3A-C).

**Figure 2.**
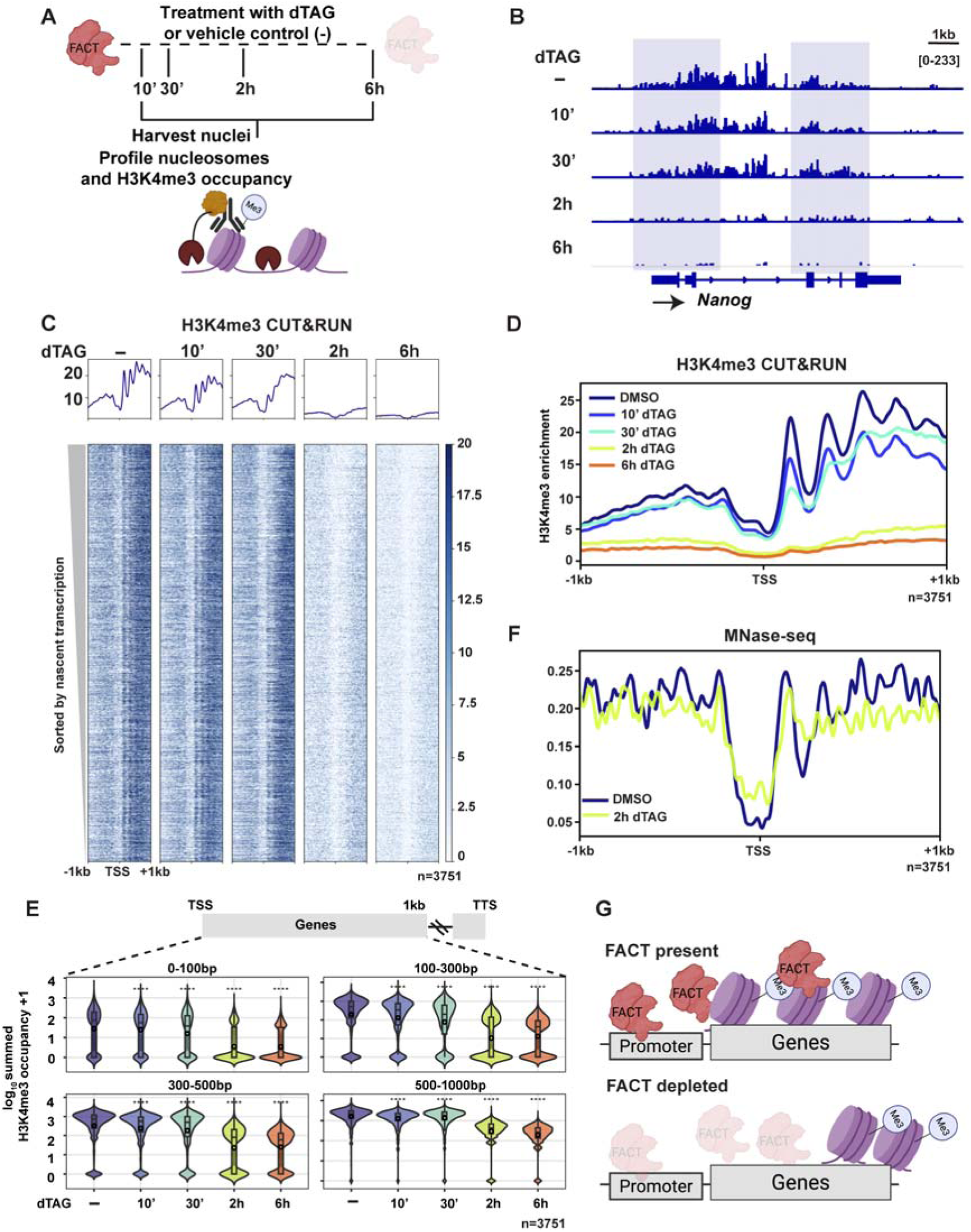
FACT depletion causes disrupted nucleosome organization and H3K4me3 occupancy at the 5’ ends of genes. **(A)** Schematic of experimental design. SPT16-dTAG ES cells were treated with dTAG-13 or vehicle control (−; DMSO) for indicated times, followed by nuclei isolation and profiling of H3K4me3 (CUT&RUN) and nucleosome occupancy (MNase-seq). **(B)** Genome browser tracks showing H3K4me3 CUT&RUN signal over the *Nanog* locus across the treatment time course in SPT16-dTAG ES cells. **(C)** Heatmap of H3K4me3 CUT&RUN signal at TSSs ± 1 kb in SPT16-dTAG ES cells across the treatment time course. Top quintile of highly transcribed genes (n = 3,751) are sorted by nascent transcription levels^83^. Metaplots (top) show average signal. **(D)** Metaplot of H3K4me3 CUT&RUN signal in SPT16-dTAG ES cells at TSSs ± 1 kb for highly transcribed genes (n = 3,751) across treatment conditions. **(E)** Violin plots showing summed H3K4me3 occupancy (log_₁₀_+ 1) across four gene body regions (0-100 bp, 100-300 bp, 300-500 bp, and 500-1000 bp downstream of TSS) following SPT16 depletion (n = 3,751 genes). ****p < 0.0001 (For each genomic window summed occupancy was compared across conditions using a Kruskal–Wallis test. Pairwise differences relative to DMSO were assessed using two-sided Wilcoxon rank-sum tests with Bonferroni correction; significance is indicated by asterisks). **(F)** Metaplot of MNase-seq signal at TSSs ± 1 kb for highly transcribed genes (n = 3,751) in vehicle control (DMSO) versus 2 h dTAG-13-treated SPT16-dTAG ES cells. **(G)** Model depicting FACT function in maintaining promoter-proximal chromatin architecture. When FACT is present (top), H3K4me3 is enriched and nucleosomes are well-positioned at the 5’ end of genes. Upon FACT depletion (bottom), H3K4me3 is lost and nucleosome organization is disrupted. See also Figures S2 and S3.

To quantify H3K4me3 loss over the depletion time course, we measured summed H3K4me3 occupancy over different bins within 1 kb downstream of the TSS and observed that the reduction in occupancy was greatest within the first 300 bp, though significant loss was detected across all bins examined (Fig. 2E). Notably, H3K4me3 loss at 10 minutes of SPT16 depletion preceded the onset of TF mislocalization, which was first detected at 30 minutes post depletion (Fig. 1G-H), suggesting that chromatin changes are a prerequisite for aberrant TF occupancy.

We then assessed nucleosome positioning. Within our H3K4me3 CUT&RUN data, nucleosome positions were clearly detectable, and we observed a loss of aggregate nucleosome phasing starting at 30 minutes of SPT16 depletion and 2 hours of SSRP1 depletion (Fig. 2D, S2E, S3A-C). We also performed MNase-seq after 2 hours of SPT16 or SSRP1 depletion to determine nucleosome positioning. We found a shift in the +1 nucleosome away from the TSS as well as a loss of nucleosome phasing over the top quartile of highly transcribed genes at 2 hours of FACT depletion (Fig. 2F, S3D). Given the observed shift in the +1 nucleosome and 5’ loss of H3K4me3, we asked whether H3K4me3 modified nucleosomes were being pushed into the gene body, causing a redistribution away from 5’ ends. To test this, we called H3K4me3 peaks in the time course depletion datasets and assigned genic peaks to bins across the gene body (Fig. S3E). Peaks within the first 1 kb of the gene show a progressive decrease, while peaks within 1-3 kb and 3-5 kb show a progressive increase over depletion time, supporting a 3’ shift of H3K4me3 modified nucleosomes.

To assess whether loss of H3K4me3 and nucleosome phasing were a specific consequence of FACT depletion rather than a general perturbation of the pluripotency network in ES cells, we obtained SOX2-dTAG and NANOG-dTAG cell lines and assessed H3K4me3 occupancy after TF depletion^48^. After 2 hours of either SOX2 or NANOG depletion, we did not observe any changes to H3K4me3 or nucleosome phasing, suggesting that these chromatin alterations are unique to FACT loss (Fig. S3F-H). Together, these data indicate that FACT is required to maintain H3K4me3 occupancy and nucleosome organization at the 5’ ends of actively transcribed genes, in agreement with previous studies in *S. cerevisiae* and *D. melanogaster*^28,31^. The temporal precedence of these chromatin defects relative to TF mislocalization suggests that FACT-dependent chromatin architecture normally restricts TF access to gene bodies (Fig. 2G).

### Single molecule sequencing confirms nucleosome loss over the 5’ ends of gene bodies upon FACT depletion

Our CUT&RUN and MNase-seq analyses provided insight into the genomic landscape upon FACT depletion. However, to overcome the limitations of using short-read sequencing to observe bulk averages at discrete loci, we employed the long-read sequencing technique Fiber-seq. This single-molecule approach enables nucleotide-resolution readout of chromatin features over multi-kilobase DNA fibers^49,50^. The method relies on methylation of adenines within unprotected DNA regions, followed by genomic DNA isolation and long-read sequencing.

We depleted SPT16 or SSRP1 for 2 hours, treated nuclei with Hia5 methyltransferase, and isolated ∼18 kb gDNA fragments. Libraries were subjected to single-molecule circular consensus sequencing on a PacBio REVIO instrument. All samples yielded coverage between 20-40X and an average fiber length of ∼10 kb. Sites with m6A were predicted, and nucleosomes were called by identifying stretches of DNA protected from methylation (Fig. 3A).

**Figure 3.**
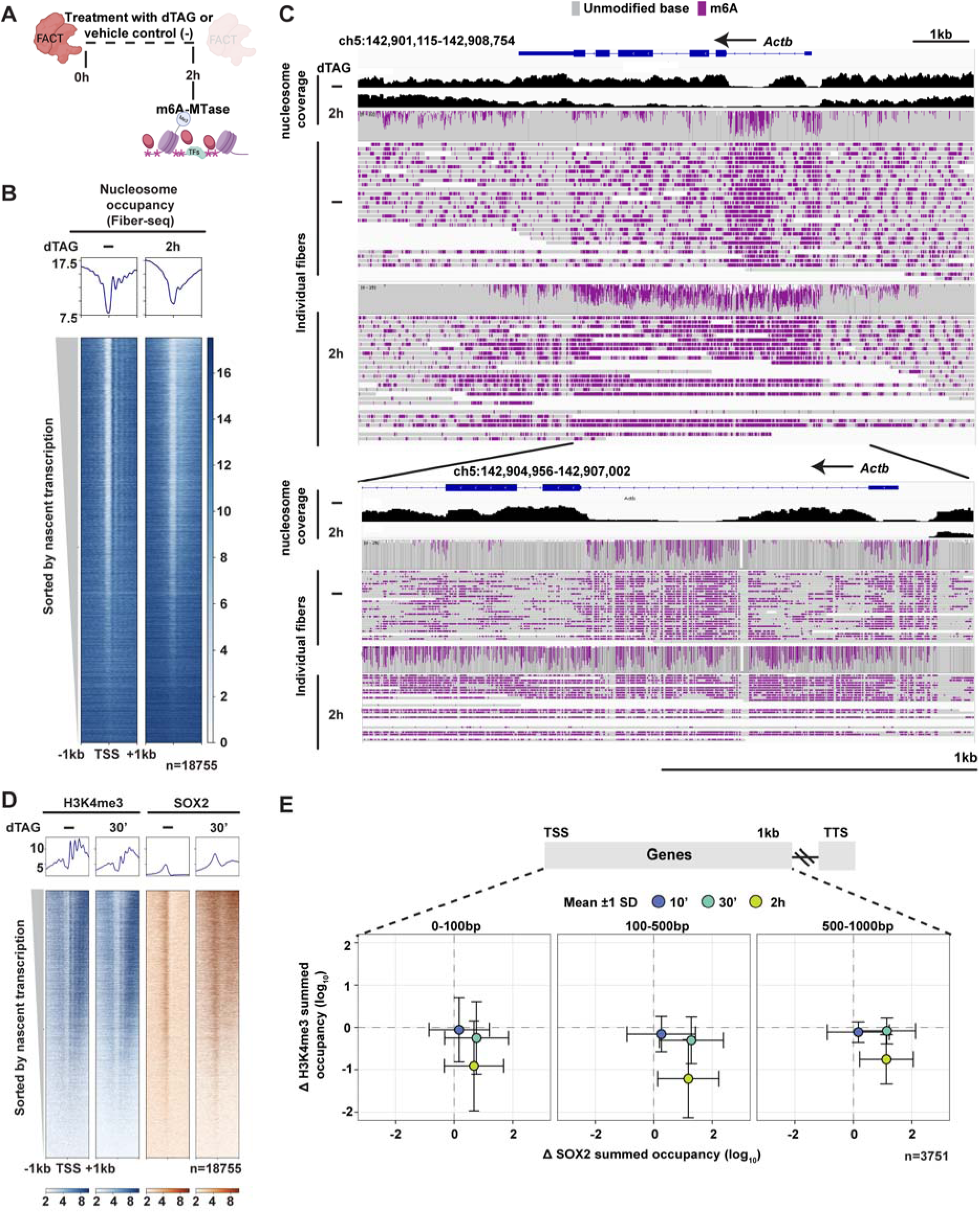
H3K4me3 loss and transcription factor accumulation occur at the same genomic locations. **(A)** Schematic of Fiber-seq experimental design. SPT16-dTAG ES cells were treated with dTAG-13 or vehicle control (−; DMSO) for 2 h, followed by nuclei isolation and treatment with DNA adenine methyltransferase, Hia5. Genomic fragments were subjected to PacBio long-read sequencing. **(B)** Heatmap of nucleosome occupancy derived from Fiber-seq analysis at TSS ± 1 kb in SPT16-dTAG ES cells comparing vehicle control (−; DMSO) and 2 h dTAG-13 treatment. Genes (n = 18,755) are sorted by nascent transcription levels^83^. Metaplot (top) shows average nucleosome occupancy. **(C)** Chromatin fibers from Fiber-seq over *Actb* locus. Top panels show a broad view (chr5:142,901,115-142,908,754); bottom panels show a zoomed in view (chr5:142,904,956-142,907,002). Nucleosome coverage tracks (top of each section) and individual fiber representations are shown for vehicle control (−; DMSO) and 2 h dTAG-13 treatment. Gray indicates unmodified bases; purple indicates N6-methyladenine (m6A) marks generated by Hia5 methyltransferase, which labels open DNA. **(D)** Side-by-side heatmaps of H3K4me3 (left) and SOX2 (right) CUT&RUN signal at TSSs ± 1 kb in SPT16-dTAG ES cells comparing vehicle control (−; DMSO) and 30 min dTAG-13 treatment. Genes (n = 18,755) are sorted by nascent transcription levels^83^. Metaplots (top) show average signal across all genes. **(E)** Scatterplots comparing changes in H3K4me3 occupancy (Δ K4me3, relative to vehicle control, y-axis) versus changes in SOX2 occupancy (Δ SOX2, relative to vehicle control, x-axis) across three gene body regions (0-100 bp, 100-500 bp, and 500-1000 bp downstream of TSS) at 10 min (blue), 30 min (teal), and 2 h (yellow) of SPT16 depletion (n = 3,751 genes). Data points represent mean ± 1 SD. See also Figure S4.

Genome-wide visualization of m6A signal revealed interrupted m6A peaks resembling nucleosome phasing in vehicle control-treated samples. This pattern was lost in FACT-depleted samples, with no increase in accessibility genome-wide (Fig. S4A). Visualization of nucleosomes over promoters sorted by nascent transcription showed loss of nucleosomes and phasing over highly transcribed genes after 2 hours of SPT16 depletion or SSRP1 depletion (Fig. 3B, Fig. S4B). We then visualized m6A coverage in untreated and FACT-depleted samples over *Actb*, a housekeeping and highly transcribed gene. In vehicle control-treated samples, we observed a cluster of m6A-marked bases coinciding with absence of nucleosomes upstream of the gene, with regularly interspersed m6A calls throughout the gene, indicative of nucleosomes across the gene body (Fig. 3C, S4C). However, when SPT16 or SSRP1 were depleted, we observed numerous clusters of m6A-modified bases upstream of the gene and along the gene body. This coincided with absence of nucleosome calls over the gene body, with a pileup of nucleosome calls downstream of the gene (Fig. 3C, S4C). Supporting our observation of H3K4me3 peaks further into the gene body compared to controls (Fig. S3E), pile up of nucleosome calls observed towards the end of the gene suggests that nucleosomes are being pushed along the gene body. Together, these results corroborated our findings from short-read sequencing methods and underscore FACT’s role in maintaining nucleosome occupancy and organization across the genome.

### Coincident nucleosome loss and SOX2 accumulation over the 5’ ends of gene bodies upon FACT depletion

After establishing that FACT depletion leads to a unique disruption of ES cell chromatin architecture, we sought to identify whether changes in histone modifications, nucleosomes, and TFs occur over the same genomic locations. We hypothesized that TF mislocalization over gene bodies would be most severe at locations where chromatin architecture was severely disrupted due to loss of nucleosomes. To assess this, we plotted SOX2 and H3K4me3 CUT&RUN datasets together, sorted by nascent transcription. As predicted, we qualitatively observed increased SOX2 occupancy over gene bodies where H3K4me3 was most severely reduced from the 5’ ends of gene bodies in both SPT16 and SSRP1 depletion (Fig. 3D, S4D).

To quantify this change, we plotted the mean change in H3K4me3 and SOX2 occupancy compared to vehicle control-treated samples over highly transcribed genes (Fig. 3E). We observed a shift in mean occupancy toward increase in SOX2 and decrease in H3K4me3 occupancy in cells depleted of SPT16. This trend progressively intensified over the depletion time course with greater mean SOX2 accumulation as SPT16 was depleted for 2 hours (Fig. 3E, S4E). SSRP1 depletion showed a similar, albeit less pronounced, trend (Fig. S4F). Together, these results support that changes in chromatin and TF occupancy are occurring at the same genes.

### Acute FACT depletion causes partially irreversible changes in the epigenomic landscape

Given that FACT loss results in disruptions to the ES cell epigenomic landscape, we asked whether restoration of FACT protein permits phenotypic recovery and epigenome restoration. To address this, we attempted to leverage the reversibility of the dTAG system by washing out the dTAG drug from media to permit protein restoration as reported previously for other proteins^51^. We depleted SPT16 or SSRP1 for 2 hours and washed out dTAG for 4 or 22 hours (Fig. 4A). Compared to vehicle control-treated cells, we did not observe protein restoration of either SPT16 or SSRP1 after 4 hours of dTAG washout (Fig. 4B-C, S5A-B). Washout of dTAG into vehicle control containing media for 22 hours resulted in severe loss of cell viability likely due to lack of protein restoration (Fig. 4D-E, S5C-D).

**Figure 4.**
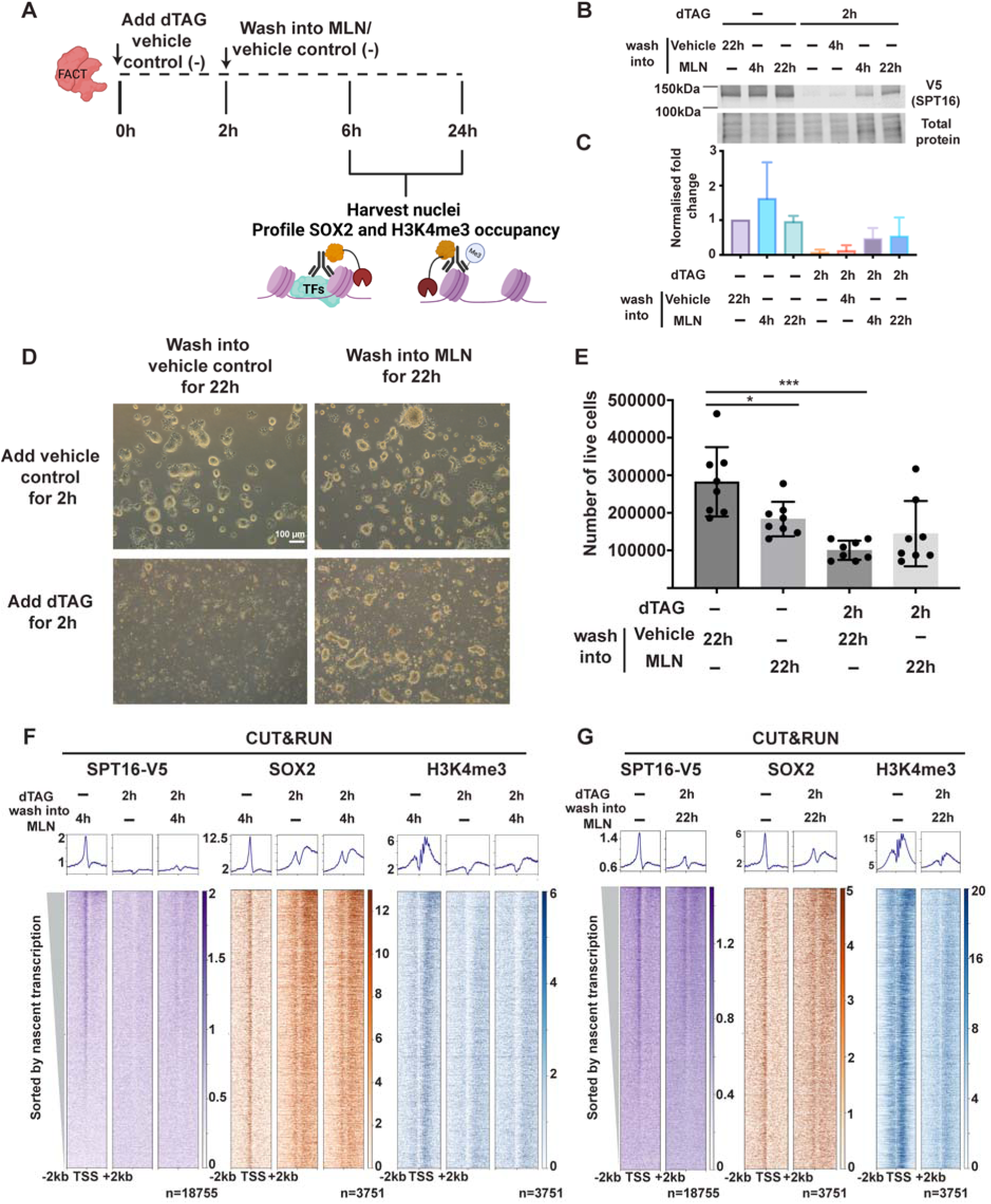
Restoration of FACT protein after degradation partially rescues transcription factor mislocalization and H3K4me3 loss. **(A)** Schematic of experimental design for FACT rescue experiments. SPT16-dTAG ES cells were treated with dTAG-13 or vehicle control for 2 h, followed by washout into MLN4924 or vehicle control. Cells were harvested at 4 or 22 h post-MLN treatment for analysis of SOX2 occupancy and H3K4me3. **(B)** Western blot analysis of SPT16-V5 protein levels across treatment conditions. **(C)** Quantification of protein levels, normalized to total protein. Data represent normalized fold change ± SD. **(D)** Representative brightfield microscopy images of SPT16-dTAG ES cells after 22 h of washout into vehicle control (left) or MLN4924 (right), following initial 2 h treatment with vehicle control (top) or dTAG-13 (bottom). **(E)** Quantification of cell viability across treatment conditions. Data represent mean ± SEM (n ≥ 6 biological replicates). *p < 0.05, ***p < 0.001 (One way ANOVA, Kruskal-Wallis test for multiple comparisons). **(F)** Heatmaps of SPT16-V5 (left), SOX2 (middle), and H3K4me3 (right) CUT&RUN signal at TSSs ± 2 kb in SPT16-dTAG ES cells across treatment conditions: 2 h vehicle control followed by 4 h MLN4924, 2 h dTAG followed by 4 h vehicle control, and 2 h dTAG treatment followed by 4 h MLN4924 treatment. Highly transcribed genes (n = 3,751) are sorted by nascent transcription levels^83^. Metaplots (top) show average signal. **(G)** Heatmaps of SPT16-V5 (left), SOX2 (middle), and H3K4me3 (right) CUT&RUN signal at TSSs ± 2 kb in SPT16-dTAG ES cells across treatment conditions: 2 h vehicle control followed by 22 h vehicle control, and 2 h dTAG treatment followed by 22 h MLN4924 treatment. Highly transcribed genes (n = 3,751) are sorted by nascent transcription levels. Metaplots (top) show average signal. See also Figure S5.

To enhance protein recovery, we performed the washout of dTAG into media containing MLN4924 (hereafter referred to as MLN; Fig. 4A). MLN is a potent inhibitor of NEDD8-activating enzyme, which in turn inactivates CRL E3 ubiquitin ligases, preventing polyubiquitylation of FKBP12 (F36V) tagged proteins^52^. Washout of dTAG into media containing MLN allowed for partial restoration of SPT16 and SSRP1 protein levels (Fig. 4B-C, S5A-B). We note that SPT16 and SSRP1 protein levels reached ∼60% and ∼30% of control treated levels, respectively, by 22 hours of washout into media containing MLN (Fig. 4B-C, S5A-B). Phenotypic recovery was only partially achieved in cells acutely depleted of SPT16 and allowed to recover in the presence of MLN (Fig. 4D-E). Cells acutely depleted of SSRP1 and washed into MLN were able to recover phenotypically to levels qualitatively comparable to vehicle control-treated cells receiving MLN (Fig. S5C-D). These data further suggest distinct roles of the two FACT subunits in ES cells.

Next, we asked whether the epigenomic landscape reflected these phenotypic observations. We profiled occupancy of SOX2 and H3K4me3 after 2 hours dTAG treatment following 4 or 22 hours washout into MLN-containing media. First, we noted that FACT occupancy was not restored in the 4 hour washout condition (Fig. 4F). Consistent with this, we observed mislocalized occupancy of SOX2 and continued 5’ loss of H3K4me3 over actively transcribed genes (Fig. 4F, S5E). By 22 hours of dTAG washout into MLN, we observed only partial recovery of the epigenomic landscape, with persistent gene body SOX2 occupancy and 5’ loss of H3K4me3, especially over highly transcribed genes in both SPT16 and SSRP1 acute depletions (Fig. 4G, S5E,F).

We hypothesized that longer FACT depletions would cause progressive disruptions from which recovery would not be possible. To test this, we assessed whether cells could phenotypically recover following longer FACT depletions followed by washout into MLN. We observed that cells depleted of SPT16 for 4 or 6 hours did not phenotypically recover even with addition of MLN to support protein recovery (Fig. S5G). Together, these data demonstrate that even transient FACT depletion is sufficient to induce persistent epigenomic alterations and compromise ES cell identity, suggesting that FACT-dependent chromatin organization, once disrupted, cannot be fully restored.

### FACT maintains chromatin architecture in a transcription-dependent manner

Having established that highly transcribed genes are most severely affected by FACT depletion, we asked whether these effects depend on ongoing transcription. To that end, we inhibited transcription using DRB, a selective and reversible inhibitor of RNAPII elongation that blocks CDK9-mediated phosphorylation of the RNAPII C-terminal domain (CTD; Fig. 5A). We noted a modest reduction in SPT16 occupancy upon treatment with DRB (Fig. 5B). This observation is consistent with transcription-dependent FACT recruitment. H3K4me3-marked nucleosome positions remained unchanged upon DRB inhibition for 2 hours, with a modest reduction in occupancy, likely reflecting reduced transcription (Fig. 5C-D). Dual treatment with dTAG and DRB for 2 hours showed a reduction in H3K4me3 occupancy but restoration of nucleosome phasing (Fig 5B-D, S6A-C). These data demonstrate that loss of nucleosome phasing in the absence of FACT depends on ongoing transcription, consistent with studies performed in *S. cerevisiae*^28^. Thus, FACT is dispensable for maintaining nucleosome organization when transcription is inhibited.

**Figure 5.**
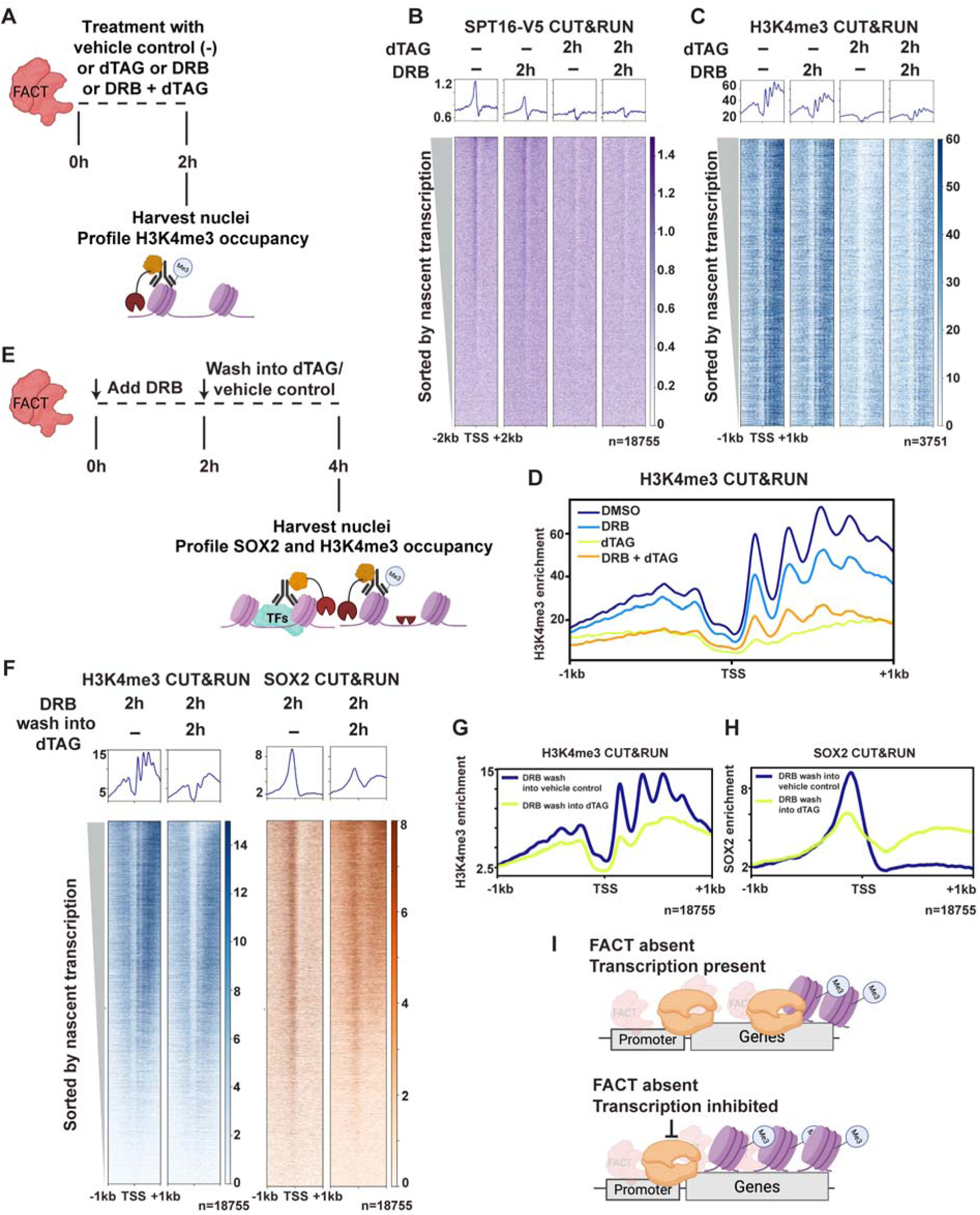
FACT-dependent chromatin architecture requires active transcription. (**A**) Schematic of experimental design to test transcription-dependence of FACT-mediated chromatin maintenance. SPT16-dTAG ES cells were treated with vehicle control (−; DMSO), DRB alone, dTAG-13 alone, or DRB combined with dTAG-13 for 2h, followed by nuclei harvest and CUT&RUN profiling of H3K4me3. (**B**) Heatmaps of SPT16-V5 CUT&RUN (TSSs ± 2 kb, n = 18,755 genes) across treatment conditions in SPT16-dTAG ES cells. Genes are sorted by nascent transcription levels^83^. Metaplots (top) show average signal. (**C**) Heatmaps of H3K4me3 CUT&RUN (TSSs ± 1 kb, n = 3,751 highly transcribed genes) across treatment conditions in SPT16-dTAG ES cells. Genes are sorted by nascent transcription levels^83^. Metaplots (top) show average signal. (**D**) Metaplot of H3K4me3 CUT&RUN signal at TSSs ± 1 kb across all four treatment conditions (DMSO, DRB alone, dTAG alone, DRB + dTAG) in SPT16-dTAG ES cells. (**E**) Schematic of experimental design to test whether FACT is required for chromatin restoration after transcription resumption. SPT16-dTAG ES cells were first treated with DRB for 2 h to inhibit elongation, then washed into dTAG-13 (to deplete FACT) or vehicle control for an additional 2 h before nuclei harvest. (**F**) Heatmaps of H3K4me3 (left) and SOX2 (right) CUT&RUN signal at TSSs ± 1 kb (n = 18,755 genes) across treatment conditions in SPT16-dTAG ES cells. Genes are sorted by nascent transcription levels^83^. Metaplots (top) show average signal. (**G**) Metaplot of H3K4me3 CUT&RUN signal at TSSs ± 1 kb comparing DRB washout into vehicle control (−; DMSO) versus DRB washout into dTAG-13 in SPT16-dTAG ES cells. (**H**) Metaplot of SOX2 CUT&RUN signal at TSSs ± 1 kb comparing DRB washout into vehicle control versus DRB washout into dTAG-13 in SPT16-dTAG ES cells. (**I**) Model depicting that FACT activity requires transcription. When FACT is absent (top), nucleosome organization is disrupted during transcription. Upon FACT depletion and transcription inhibition (bottom), H3K4me3 and nucleosomes are not disrupted. See also Figure S5 and Figure S6.

Next, we asked whether FACT is required to maintain the chromatin landscape when transcription resumes after inhibition. To test this, we treated ES cells with DRB for 2 hours and washed out DRB into media containing vehicle control or dTAG to deplete SPT16 or SSRP1 (Fig. 5E). We profiled SOX2 and H3K4me3 occupancy and found that cells washed into dTAG had increased gene body SOX2 occupancy and loss of 5’ H3K4me3 and nucleosome phasing (Fig. 5F-H, S6D-F). Together, these results demonstrate that FACT is essential for maintaining chromatin architecture during transcription restoration. These findings support a model in which transcription-induced nucleosome displacement creates a continuous requirement for FACT-mediated nucleosome reassembly and organization (Fig. 5I).

### FACT depletion results in RNAPII accumulation at the 5’ ends of genes

Given the transcription-dependent chromatin changes observed following rapid FACT depletion, we hypothesized that these changes could lead to 5’ accumulation of RNAPII and decreased ability for RNAPII to successfully elongate. To test this, we profiled RNAPII CTD serine 5 phosphorylation (RNAPII-S5p) occupancy after depleting SPT16 or SSRP1 (Fig. 6A) and observed an accumulation in occupancy over the 5’ ends of gene bodies (Fig. 6B, S6G). Notably, this is not due to a change in total protein levels of RNAPII-S5p (Fig. SH-I). We quantified summed RNAPII-S5p chromatin occupancy within the first 1 kb of the gene body and observed an increase over gene bodies starting at 30 minutes of SPT16 depletion and 2 hours of SSRP1 depletion (Fig. 6C, S6J).

**Figure 6.**
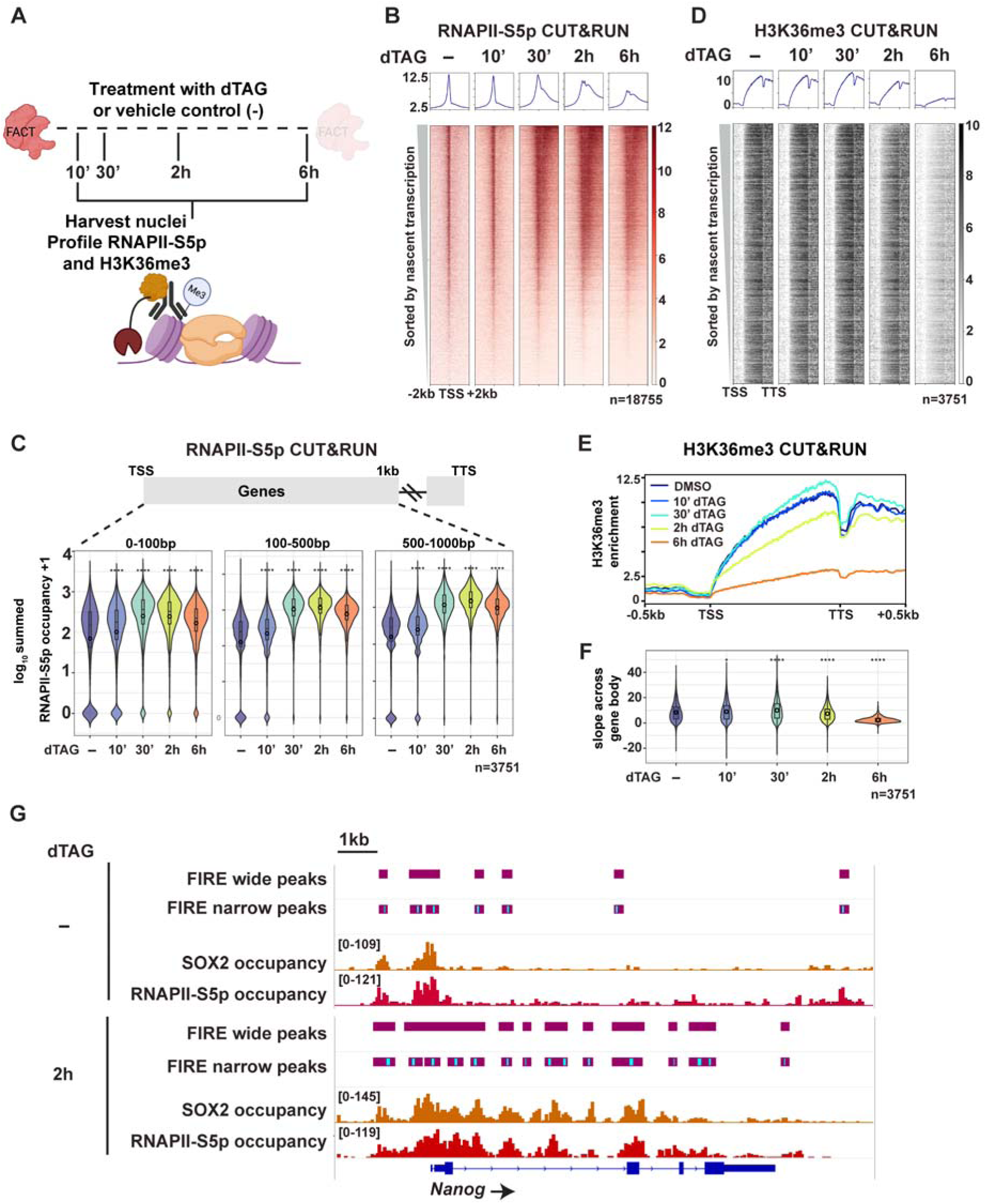
FACT depletion causes RNAPII accumulation at 5’ ends of genes and altered H3K36me3, indicative of elongation defects. **(A)** Schematic of experimental design. SPT16-dTAG ES cells were treated with dTAG-13 or vehicle control (−; DMSO) for indicated times, followed by nuclei isolation and CUT&RUN profiling of RNAPII-S5p and H3K36me3. **(B)** Heatmap of RNAPII-S5p CUT&RUN signal at TSSs ± 2 kb in SPT16-dTAG ES cells across the dTAG-13 treatment time course. Genes (n = 18,755) are sorted by nascent transcription levels^83^. Metaplot (top) shows average signal. **(C)** Violin plots showing summed RNAPII-S5p occupancy (log_₁₀_+ 1) across three gene body regions (0-100 bp, 100-500 bp, and 500-1000 bp downstream of TSS) following SPT16 depletion for top quintile of highly transcribed genes (n = 3,751 genes). *p < 0.05, ****p < 0.0001 (For each genomic window summed occupancy was compared across conditions using a Kruskal–Wallis test. Pairwise differences relative to DMSO were assessed using two-sided Wilcoxon rank-sum tests with Bonferroni correction; significance is indicated by asterisks). **D)** Metagene heatmap of H3K36me3 CUT&RUN signal across gene bodies (TSS to TTS) in SPT16-dTAG ES cells across the dTAG-13 treatment time course. Top quintile of highly transcribed genes (n = 3,751) are sorted by nascent transcription levels^83^. Metaplot (top) shows average signal. **(E)** Metaplot of H3K36me3 CUT&RUN enrichment across scaled gene bodies (TSS to TTS ± 0.5 kb) for highly transcribed genes (n = 3,751) across treatment conditions. **(F)** Violin plots quantifying the slope of H3K36me3 signal across gene bodies (n = 3,751 genes) following SPT16-dTAG depletion. ****p < 0.0001 (For each genomic window summed occupancy was compared across conditions using a Kruskal–Wallis test. Pairwise differences relative to DMSO were assessed using two-sided Wilcoxon rank-sum tests with Bonferroni correction; significance is indicated by asterisks). **(G)** Fiber-seq Inferred Regulatory Elements (FIRE) peaks over the *Nanog* locus in SPT16-dTAG ES cells treated with vehicle control (−; DMSO, top) or dTAG-13 for 2 h (bottom). Tracks show (from top to bottom): FIRE-called wide and narrow peaks, SOX2 CUT&RUN, RNAPII-S5p CUT&RUN, and gene annotation. See also Figure S6 and S7.

We hypothesized that the increase in RNAPII-S5p could be explained by an RNAPII processivity and elongation defect. To test this, we first investigated RNAPII processivity using H3K36me3 occupancy as a proxy for transcription elongation. We observed a reduction in H3K36me3 enrichment over gene bodies by 2 hours of FACT depletion, indicated by loss of enrichment from TSS to TTS as well as a decreasing slope tending toward zero over the most highly transcribed genes compared to vehicle control-treated cells (Fig. 6D-F, S7A-B). Together, these data suggest that RNAPII is likely stalled at the 5’ ends of genes and unable to complete elongation.

We then applied FIRE (Fiber-seq Inferred Regulatory Elements), a recently developed machine learning classifier that enables precise delineation of chromatin architecture, to our Fiber-seq data^53^. FIRE identifies contiguous regions of protection along single DNA molecules and classifies them by width. In vehicle control-treated samples, we observed sharp narrow peaks marking precise, reproducible accessibility at a fixed position across many fibers at the 5’ end of *Nanog*. Narrow FIRE peaks are consistent with stable positionally constrained TF occupancy and correspond to putative TF footprints^54^. Wide peaks reflect accessibility spread over a region with variable boundaries across fibers^54^. We observed these peaks at 5’ ends and carefully distributed intervals through *Nanog* (Fig. 6G). These peaks are proposed to be consistent with regionally distributed accessibility often observed at RNAPII engaged promoters^53^. Positions of these peaks aligned with peaks called from SOX2 and RNAPII-S5p CUT&RUN data from control samples (Fig. 6G). When SPT16 or SSRP1 was depleted for 2 hours, we observed an increase in both narrow and wide accessibility peaks coinciding with expanded SOX2 and RNAPII-S5p across the locus (Fig. 6G, S7C). These single-molecule data corroborate our bulk CUT&RUN findings and suggests that TF and RNAPII accumulation occurs on the same DNA molecules.

### Chromatin architectural changes precede transcriptional dysfunction caused by FACT depletion

The preceding experiments establish a clear temporal hierarchy of molecular events following FACT depletion. H3K4me3 and inferred nucleosome loss was detectable as early as 10 minutes post-depletion of SPT16 (Fig. 2C-E), followed by RNAPII accumulation and TF mislocalization at 5’ ends of genes by 30 minutes of SPT16 depletion (Fig. 1G-H, Fig. 6B, D). SSRP1 depletion led to the same cascade of events, but with a delayed onset relative to SPT16 depletion. This sequence of events raised the question of whether these chromatin perturbations ultimately impact transcriptional output. To test this, we performed transient transcriptome sequencing (TT-seq) to measure nascent transcription after 10 minutes, 30 minutes, and 2 hours of SPT16 and SSRP1 depletion (Fig. 7A). Interestingly, we observed modest or no transcriptional changes at 10 and 30 minutes of SPT16 or SSRP1 depletion (Fig. 7B-C, Fig. S7D). However, by 2 hours of FACT depletion, we observed widespread transcriptional dysregulation, with the majority of genes showing reduced transcription (Fig. 7D, Fig. S7D).

**Figure 7.**
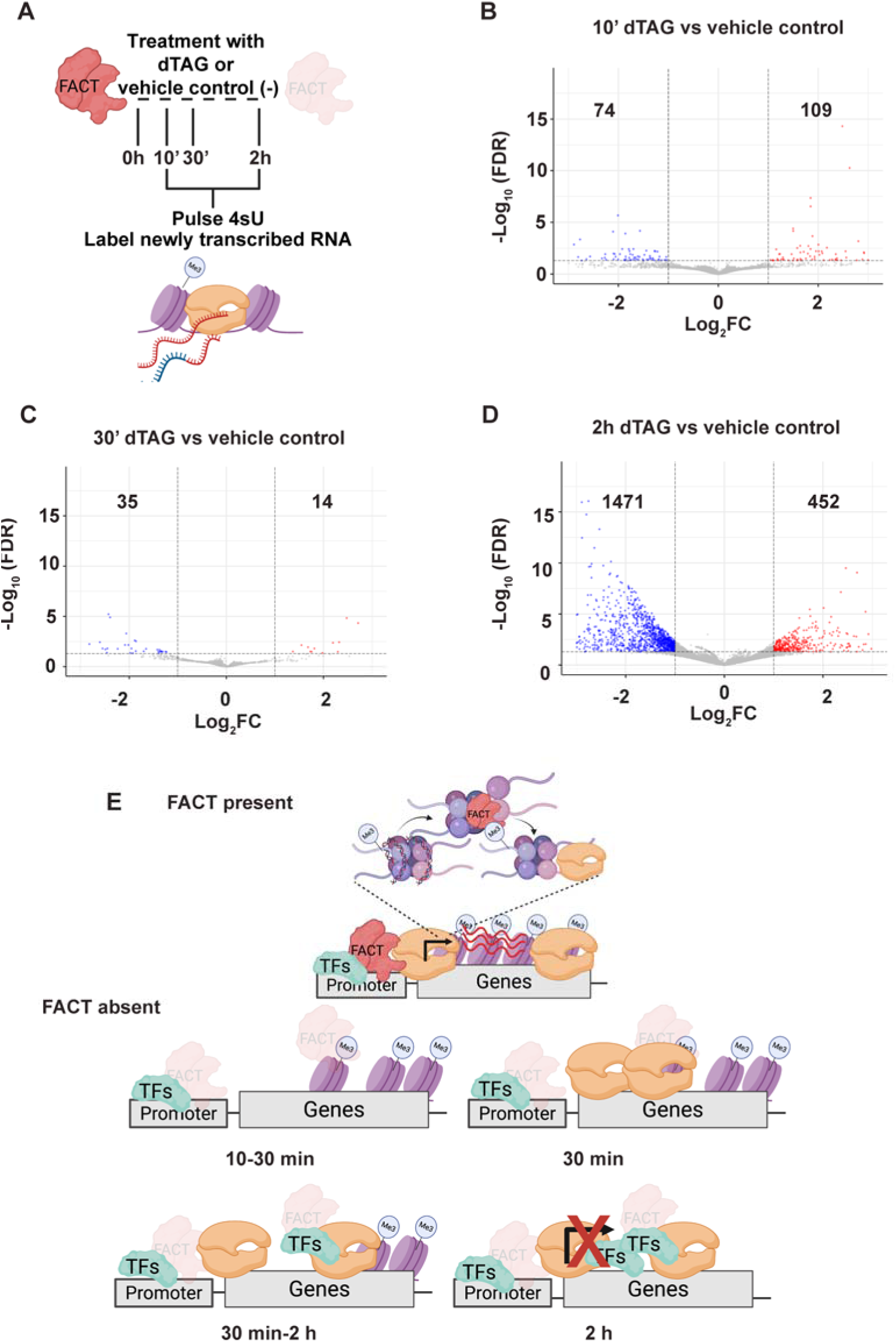
Transcription dysregulation occurs after epigenomic changes upon FACT depletion. **(A)** Schematic of experimental design for nascent transcription analysis. SPT16-dTAG ES cells were treated with dTAG-13 or vehicle control (−; DMSO) for indicated times, followed by pulse labeling with 4-thiouridine (4sU) to capture newly transcribed RNA for TT-seq analysis. **(B)** Volcano plot showing differential nascent transcription at 10 min of SPT16 depletion compared to vehicle control. X-axis shows log_₂_ fold change; y-axis shows −log_₁₀_FDR. Significantly upregulated genes (red, FDR<0.05 and Log_2_FC>1) and downregulated genes (blue, FDR<0.05 and Log_2_FC<1) are highlighted. **(C)** Volcano plot as in (B) showing differential nascent transcription at 30 min of SPT16 depletion compared to vehicle control. **(D)** Volcano plot as in (B) showing differential nascent transcription at 2 h of SPT16 depletion compared to vehicle control. **(E)** Model summarizing the temporal sequence of events following FACT depletion. FACT loss causes H3K4me3 and nucleosome loss at 5’ ends of genes. TFs are mislocalized to gene bodies, RNAPII accumulates over 5’ ends of genes resulting in eventual downregulation of transcription. See also Figure S7.

To correlate whether the transcription and chromatin changes are occurring at the same locations, we visualized H3K4me3 occupancy over significantly up and downregulated genes calculated from our TT-seq experiments. We observed reduced H3K4me3 occupancy and nucleosome phasing at 30 minutes of SPT16 depletion over genes that were downregulated by 2 hours of SPT16 depletion (Fig. S7E). H3K4me3 occupancy was more modestly disrupted over locations that were upregulated at 2 hours of SPT16 depletion (Fig S7E). This temporal delay between chromatin perturbation and transcription downregulation is consistent with our model of defective RNAPII progression and reduced H3K36me3 deposition.

During transcription, non-canonical nucleosome structures referred to as overlapping dinucleosomes (OLDNs, also known as hexasome-nucleosomes) are formed when one nucleosome collides with an adjacent nucleosome and a single H2A/H2B dimer is released. Both nucleosome remodelers and FACT have been proposed to help resolve these structures^54–58^. Therefore, we hypothesized that without FACT to resolve these structures, one reason for RNAPII stalling in the 5’ ends of genes may be the accumulation of OLDNs. Using size-selected MNase-seq to enrich for DNA protected by OLDNs, we observed an increase in OLDN occupancy over highly transcribed genes following 2 hour SSRP1 depletion (Fig. S7F). These data support a role for FACT in resolving OLDNs and suggest that altered chromatin architecture is contributing to reduced RNAPII processivity.

Together, these data reveal a temporal cascade in which FACT depletion causes 5’ loss of nucleosomes and their associated histone modifications (10-30 minutes). This chromatin disruption (loss of nucleosomes and H3K4me3) occurs prior to transcriptional inhibition, arguing transcription without FACT disrupts transcription-coupled nucleosome reassembly and, as a consequence, nucleosomes and histone modifications in 5’ regions are lost (30 minutes - 2 hours). RNAPII accumulates in 5’ regions and eventually transcription shuts down, likely reflecting FACT is required for efficient RNAPII progression through a nucleosome template *in vitro* and in vivo. Notably our 2 hour Fiber-seq data demonstrate accumulated nucleosomes that have pushed 3’, which supports reduced nucleosome disassembly due to FACT loss. In parallel, open chromatin promotes promiscuous TF binding (30 minutes - 2 hours). The delayed onset of transcriptional changes relative to chromatin perturbations demonstrates that FACT’s histone chaperone activity is a prerequisite for productive transcription (Fig. 7E).

## DISCUSSION

The precise mechanisms by which FACT maintains chromatin integrity and the consequences of its loss have remained incompletely understood. Here, using rapid depletion of FACT subunits combined with high-resolution temporal profiling, we reveal a cascade of chromatin disruption events that culminate in TF mislocalization, transcriptional collapse, and cell death. Our data support a model wherein loss of FACT prevents transcription-coupled nucleosome reassembly behind RNAPII. In addition, RNAPII accumulates in 5’ regions with higher density of nucleosomes in 3’ regions, consistent with impaired elongation due to lack of nucleosome disassembly, resulting in eventual transcriptional collapse. These data position FACT as a guardian of chromatin architecture whose loss triggers a progressive deterioration of the regulatory landscape.

### A temporal model for FACT-dependent chromatin maintenance

Our findings support FACT’s known function to reassemble nucleosomes in the wake of elongating RNAPII^18,19,21,59,60^. Upon FACT depletion, we observe a rapid sequence of events: loss of nucleosome phasing and H3K4me3 at the 5’ ends of genes, accumulation of RNAPII-S5p, progressive loss of H3K36me3 especially at the 3’ of genes, and invasion of TFs into gene bodies. Strikingly, these chromatin changes occur within 30 minutes of FACT depletion, whereas significant transcriptional downregulation is only detected 2 hours post FACT depletion.

We propose that in the absence of FACT, elongating RNAPII together with other factors, such as nucleosome remodelers, displace nucleosomes but these cannot be efficiently reassembled, leading to progressive chromatin disruption at the 5’ ends of genes. Notably, our data reveal that this disruption occurs in stages: nucleosomes first lose their precise phasing, then are pushed into the gene-body before eventual loss of the nucleosome altogether. The accumulation of RNAPII-S5p at promoter-proximal regions suggests that RNAPII progression is impaired, potentially due to collisions with improperly positioned nucleosomes or increased non-canonical nucleosome structures such as OLDNs. Notably, the temporal sequence we observe (H3K4me3 and nucleosome loss preceding RNAPII-S5p accumulation) is consistent with recent findings that H3K4me3 is required for efficient RNAPII pause-release^61,62^. H3K4me3 is subject to rapid turnover that requires continuous re-establishment by COMPASS^63^. In addition, COMPASS recruitment depends on both interaction with elongating RNAPII and recognition of existing H3K4me3^64,65^. Structural studies reveal COMPASS makes extensive contacts with nucleosomal DNA at three distinct locations that properly position its catalytic domain^66,67^. Therefore, this read-write mechanism could become compromised once H3K4me3 is displaced from its proper location due to FACT no longer being present to reassemble the nucleosome. Recent data showing FACT’s role in suppressing HK36me3 deposition on the downstream nucleosome, ensuring that deposition occurs behind the transcribing RNAPII on the upstream nucleosome, suggests that our observed reduction of H3K36me3 over the 3’ ends of genes is due to reduced nucleosome reassembly when FACT is depleted^68,69^. Therefore, aberrantly positioned nucleosomes may be suboptimal substrates for methylation. Together, these data suggest that FACT-mediated nucleosome reassembly is essential not only for maintaining nucleosome positioning but also for preserving the chromatin context required for H3K4me3 and H3K36me3 maintenance.

### Transcription factor invasion reflects chromatin accessibility, not active recruitment

The redistribution of SOX2, OCT4, and NANOG into gene bodies (Fig. 1) likely reflects a loss of physical barriers rather than a change in TF recruitment. Our data support a model wherein transcription-induced clearing of nucleosomes without FACT-mediated reassembly creates an accessible DNA landscape that TFs can occupy. This is in line with recent results from *S. cerevisiae* where the authors individually depleted histone chaperones and observed increased Msn2 and Reb1 binding in gene bodies^70^. Our data suggest that TFs occupy accessible binding sites regardless of their sequence preferences, and whether such binding is functionally appropriate. The interpretation that TFs are simply promiscuously occupying open regions of chromatin has important implications for understanding TF dynamics. We envision that TFs are constantly sampling the genome, with preference to open chromatin regions, or when a sequence preference is found, TFs have increased residence time at that location. Therefore, due to chromatin opening at the 5’ ends of genes upon FACT depletion, we hypothesize that our bulk cell genomic experiments are capturing the increased sequence sampling of the TFs profiled. We suspect that beyond the pluripotency factors examined in this study, numerous other DNA-binding proteins similarly bind newly accessible chromatin, creating a chaotic regulatory environment incompatible with proper gene expression. Notably, a recent preprint from the Milne group demonstrates that additional DNA-binding proteins, including TBP, also invade gene bodies upon FACT depletion, supporting this model^71^.

### Consequences for higher-order chromatin organization

The disruption of local chromatin structure upon FACT depletion likely extends beyond the immediate effects on local nucleosome positioning and histone modifications. The loss of H3K4me3 and disruption of nucleosome phasing at the 5’ end of genes may propagate into alterations in enhancer-promoter contacts, topologically associating domain (TAD) boundaries, and broader nuclear organization. Initial studies examining the impact of FACT on higher-order chromatin structure suggested only a minor role^72^. More recently, using the same depletion cell line, loss of FACT was shown to perturb TAD structure and increase fuzziness of nucleosomes flanking CTCF binding sites^73^. The preprint from the Milne group demonstrates that FACT loss leads to the dissolution of nanoscale domains^71^.

### Heterogeneous cellular responses to FACT loss and recovery

Our rescue experiments using MLN4924 to restore FACT protein levels revealed heterogeneity in cellular responses to FACT depletion/restoration (Fig. 4). While a subset of cells recovered phenotypically after MLN treatment, others failed to recover. This partial rescue with acute (2 hour) depletions and lack of rescue observed with longer (4 and 6 hour) depletions, suggests the existence of a “point of no return” beyond which epigenomic damage due to FACT loss becomes irreversible. We propose that cells experiencing FACT depletion follow divergent trajectories depending on the extent of chromatin disruption at the time of FACT restoration. For example, in a two-state model, in cells where chromatin damage remains limited, restored FACT can reassemble proper nucleosome architecture, re-establish histone modification patterns, and normalize TF localization. In the second state, cells where disruption has progressed beyond a critical threshold, the accumulated damage to chromatin structure and TF localization is incompatible with recovery. These cells undergo differentiation and/or cell death, explaining both the partial rescue of viability and the incomplete normalization of bulk chromatin profiles. Single-cell approaches will be valuable for directly testing this model and identifying the molecular features that distinguish recoverable from non-recoverable cells.

### Distinct contributions of SPT16 and SSRP1 to FACT function

Throughout our experiments, we observed consistent temporal differences between SPT16 and SSRP1 depletion and resulting phenotypes. TF mislocalization and RNAPII accumulation initiated by 30 minutes of SPT16 depletion but required 2 hours following SSRP1 depletion. Relatedly, the differential recovery capacity we observed with SSRP1-depleted cells recovering more fully than SPT16-depleted cells upon dTAG washout with MLN-containing media suggests that the more rapid chromatin deterioration following SPT16 loss creates damage that is harder to reverse. Biochemical studies demonstrate that SPT16 provides the primary histone-binding surfaces of the FACT complex, directly engaging both H2A-H2B dimers and H3-H4 tetramers during nucleosome reorganization. SSRP1, while essential for FACT function, may play a more modulatory role in DNA binding and substrate recognition^12–14,18,74,75^. Therefore, our findings are consistent with SPT16 being more directly responsible for the core histone chaperone activity of FACT.

### Interplay with nucleosome remodelers

FACT does not act in isolation but rather functions within a network of nucleosome remodelers that collectively maintain chromatin integrity during transcription. Of particular relevance is the nucleosome remodeler CHD1, which has been shown to interact with FACT and to function in nucleosome reassembly during elongation^19,56,76^. CHD1 recognizes H3K4me3-marked nucleosomes and promotes their proper spacing, suggesting a functional handoff between FACT-mediated histone deposition and CHD1-mediated nucleosome positioning^77,78^. Our lab and others have investigated the role of CHD1 in ES cells, where CHD1 is required for ES cell identity and maintaining hypertranscription^79–81^. The rapid loss of H3K4me3 upon FACT depletion may therefore compound chromatin disruption by eliminating the substrate for CHD1 activity. It will be interesting to test the interplay between these two factors further in this system.

Our observation of increased OLDN accumulation at highly transcribed genes following FACT depletion provides additional mechanistic insight into this feed-forward model. Both FACT and nucleosome remodelers have been implicated in resolving these non-canonical structures^54–56,58,82^. The accumulation of OLDNs in FACT-depleted cells suggests that without FACT-mediated resolution, these structures persist and may physically impede subsequent rounds of transcription.

### Implications for disease

The rapid loss of viability in murine ES cells, beginning as early as 6 hours after SPT16 or SSRP1 depletion, underscores a critical reliance on the FACT complex to maintain the pluripotent state. ES cells have high transcriptional activity, a unique chromatin landscape, and appear particularly dependent on FACT function^13,14,36,38,39^. This sensitivity may reflect the dynamic chromatin environment of pluripotent cells, where continuous transcription of lineage-specifying genes must be balanced against maintenance of the undifferentiated state.

FACT function is also important in rapidly dividing cancer cells where FACT subunits are frequently overexpressed in tumors and have been explored as therapeutic targets^34,40–43^. Our findings provide mechanistic insight into why FACT inhibition might be selectively toxic to cancer cells: the high transcriptional demands of rapidly proliferating cells likely render them particularly sensitive to disruptions in nucleosome reassembly. The cascade of chromatin disruption we describe, from nucleosome loss to TF mislocalization to transcriptional collapse, may represent a generalizable mechanism by which FACT inhibition achieves its anti-proliferative effects.

## Conclusions

Our study establishes FACT as a critical guardian of chromatin architecture whose continuous activity is required to maintain the epigenomic landscape at transcribed genes. By revealing the temporal sequence of events following FACT depletion, we demonstrate that chromatin disruption precedes and likely causes transcriptional dysregulation, rather than simply accompanying it. This work provides a foundation for understanding how histone chaperones maintain genome function and suggests that the dynamic interplay between transcription and chromatin structure is far more delicate than previously appreciated.

### Limitations of the study

While our findings in murine ES cells provide a high-resolution temporal map, it remains to be seen if these mechanisms are conserved in other cell types. Second, while our bulk genomic approaches provided high temporal resolution, they cannot capture cell-to-cell variability in response to FACT depletion. Third, our study focused on a limited set of TFs and while we propose that other DNA-binding proteins similarly invade newly accessible chromatin, we have not directly tested this hypothesis. Finally, while we propose a model in which FACT reassembles nucleosomes behind elongating Pol II, our data are also consistent with alternative mechanisms, including roles for FACT in nucleosome stability, histone exchange, or coordination with other nucleosome remodelers such as CHD1. Biochemical reconstitution experiments and targeted perturbation of potential FACT collaborators will be essential to distinguish among these possibilities.

## METHOD DETAILS

### Cell culture

ES-E14TG2a (E14) embryonic stem cells from male *Mus musculus* origin (RRID:CVCL9108) were grown at 37°C and 5% CO_2_ in DMEM base medium (Sigma) supplemented with 1X nonessential amino acids (Gibco), 2 mM L-glutamine (Gibco, 10% FBS (Sigma), 0.129 mM β-mercaptoethanol (Sigma), 1000 U/mL LIF, 3 µM CHIR99021 GSK inhibitor and 1 µM PD0325091 MEK inhibitor. Cells were grown in feeder-free conditions on 10 cm plates gelatinized with 0.2% porcine skin gelatin type A, passaged every 48 h using 0.5% trypsin and split at a ratio of ∼1:8 with fresh medium. Routine anti-mycoplasma cleaning was conducted, and cell lines were screened using PCR to confirm absence of mycoplasma.

### Cell line generation, dTAG depletion, and other treatments

SPT16-dTAG-3XV5 cell lines were generated using CRISPR/Cas9 directed homologous recombination. sgRNAs were designed using CRISPICK and cloned into px330 plasmid containing puromycin resistance cassette (sgRNA listed in Supplementary table 1). Homology constructs with ∼500 bp of homology on either side and FKBP12(F36V) and 3XV5 were purchased from ThermoFisher (GeneArt). Low passage wild type ES cells were transfected with 3 µg px330-sgRNA, 3 µg of the homology construct and FuGene HD to generate endogenously tagged homozygous depletion cell lines. 48 h post transfection, the media was replaced with fresh media containing 2 µg/mL puromycin for 24 hours. Following transfection, single clones were picked, allowed to grow in 96 well plates and screened for successful targeting of both alleles using PCR genotyping and Sanger sequencing (oligos listed in Supplementary table 1). SSRP1-dTAG cells were generated in ES E14 cells by CRISPR-Cas9 using the SpCas9(BB)-2A-Puro (PX459) V2.0 plasmid (Addgene #62988) as previously described, targeting the SSRP1 gene at the end of the ORF and a SSRP1-linker-dTAG homology donor plasmid with homology arms amplified using oligos (listed in Supplementary table 1)^84^. Cells were transfected using Lipofectamine 3000 using 0.5 µg of the sgRNA plasmid and 2 µg of the donor plasmid. Cells were sparsely seeded on a 10 cm dish 24 h post transfection and selected with Puromycin (2 µg/mL) for 48 h. After one week in culture, individual clones were picked manually with a pipette and moved to a 96-well plate. Following transfection, single clones were picked, allowed to grow in 96 well plates and screened for successful targeting of both alleles using PCR genotyping and Sanger sequencing (oligos listed in Supplementary table 1).

Cells were depleted of SPT16 or SSRP1 protein by addition of dTAG-13 dissolved in 100% DMSO at 1 mM and added at 1 µM premixed in fresh 2i/LIF medium. Cells were incubated with either dTAG-13 or 0.1% DMSO (vehicle) for mentioned time periods to deplete FACT complex subunits. Depletion was confirmed by western blotting. Cells were cultured on 10 cm plates undisturbed for 48 h prior to dTAG depletion ensuring that relevant effects are not due to passaging related stress. DRB (Sigma Millipore; D1916) for RNAPII elongation inhibition was dissolved in DMSO at 1 mM and added at 100 µM in 2i + LIF media for specified times. MLN4924 (Cell Signaling; S5923S) for E3-ubiquitin ligase inhibition was dissolved in DMSO at 1 mM and added at 100 µM in 2i + LIF media for specified times. SOX2 and NANOG-dTAG ES cells were cultured and depleted with dTAG-13 as previously described^48^.

### Cell growth assay

Cells were plated in 6 wells at a density of 50,000 cells per well. After 6, 12, and 24 h of SPT16 or SSRP1 depletion cells were trypsinized and pipetted to a single cell suspension. Cells were then mixed with 1:1 ratio of Trypan blue and counted on a TC20 cell counter (Biorad) to distinguish live cells. Two biological replicates were conducted for each cell line. For cell growth assays following dTAG and MLN treatment, cells were counted after 6 h or 24 h of dTAG and MLN addition.

### Western blot analysis

Crude protein extractions were performed using RIPA buffer (150 mM NaCl, 1% IGEPAL CA-630, 0.5% sodium deoxycholate and 25 mM Tris-Cl, pH 7.4) with freshly added protease inhibitors and flash frozen immediately after extraction. Histone proteins were extracted using an acid-based method^79^. Cells were lysed in 200 µl Triton Extraction Buffer (TEB; 0.5% Triton X-100, 2 mM PMSF, 0.02% sodium azide) for 10 min. Lysate was centrifuged for 10 min at 4°C at 650 g and supernatant was discarded. Pellets were washed in 100 µL TEB and centrifuged as before. Supernatant was discarded and lysate was resuspended in 50 µL 0.2N HCl and rotated overnight at 4°C. Extracts were then centrifuged for 15 min at 4°C, supernatants were retained, and flash frozen. Protein concentration was measured using the Pierce BCA protein assay kit (Thermo). 30 µg of protein (10 µg for histone proteins) was diluted in RIPA buffer with 10 mM dithiothreitol (DTT) and 1X Laemmeli sample buffer before being loaded on Tris-acrylamide gels (8% acrylamide for SSRP1, SPT16, RNAPII, OCT4, SOX2, 15% acrylamide for post translational modifications) for western blot analysis. Proteins were transferred to nitrocellulose membranes at 20V overnight. Loading was measured with REVERT 700 total protein stain (LICORbio, 926-11011) and imaged by LI-COR (LI-COR Odysset DLx Imager). Membranes were blocked for 1 h in Intercept Blocking Buffer (927-60001) at room temperature. Primary antibodies were added overnight (Antibodies diluted in 1X PBST were: V5 epitope; Invitrogen 46-0705 at 1:5000, SSRP1; Biolegend 609702 at 1:500, SPT16; Cell Signaling 12191S at 1:1000, OCT4; Invitrogen 701756 at 1:1000, SOX2; Active Motif 39843 at 1:1000, NANOG; Cell Signaling 8822S at 1:1000, RNAPII-S5p; Invitrogen MA1-46093 at 1:1000, H3K4me3; Millipore 05-745R at 1:2500, H3K36me3; Invitrogen MA5-24687 at 1:1000). Following primary antibody incubation, fluorescent conjugated 800 nm secondary antibody (LICOR 926-32210, 926-32211, 1:10,000 in 1XPBST) was added for 1 h in the dark and membranes were imaged using the LI-COR. Protein quantification was conducted using ImageJ. For total protein quantification identical areas for each lane were selected, pixel intensity was measured and subtracted from a background pixel intensity. For target protein quantification, identical size areas immediately surrounding the target band were selected, pixel intensity was measure and subtracted from background pixel intensity. Target protein quantification was made relative to total protein quantification and then made relative to vehicle control treated sample. Two biological replicates were conducted per target for each cell clone of each cell line.

### CUT&RUN

CUT&RUN was performed as previously described using the following modifications^85–89^. Briefly 100,000 nuclei were isolated from mESCs or *Drosophila* S2 cells using a hypotonic buffer (20 mM HEPES-KOH pH 7.9, 10 mM KCl, 0.5mM spermidine, 0.1% Triton X-100, 20% glycerol and freshly added protease inhibitors). Nuclei were counted after isolation and combined in a 90 (ES cells) to 10 (S2 cells) ratio for spike in normalization. Nuclei were then bound to lectin coated concanavalin A magnetic beads (Polysciences, 50 µL beads per 100,000 nuclei). Immobilised nuclei were treated with blocking buffer for 5 minutes at room temperature (20 mM HEPES pH 7.5, 150 mM NaCl, 0.5 mM spermidine, 0.1%BSA, 2 mM EDTA and fresh protease inhibitors) and washed with the same buffer without EDTA (Wash buffer). Nuclei were then incubated on a rotator for 1 h at room temperature with Wash buffer containing primary antibody (V5 epitope; Invitrogen 46-0705 at 1:100, OCT4; Invitrogen 701756 at 1:100, SOX2; Active Motif 39843 at 1:100, NANOG; Active motif 9919002 at 1:100, RNAPII-S5p; Invitrogen MA1-46093 at 1:100, H3K4me3; Millipore 05-745R at 1:100, H3K36me3; Invitrogen MA5-24687 at 1:100). Controls lacking a primary antibody were subjected to same conditions but incubated with IgG Rabbit (1:500, Abcam ab37415) in Wash buffer. Beads bound by nuclei and primary antibody were then washed twice with Wash buffer and then resuspended in Wash buffer containing pA-MNase (for rabbit antibodies) or pAG-MNase (for mouse antibodies). Samples were rotated for 30 minutes at room temperature and washed twice with Wash buffer following binding of pA/pAG MNase. Samples were then pre-equilibrated to 0°C in an ice water bath for 5 min and 3 mM CaCl_2_ was added to activate pA/pAG MNase cleavage. After digestion for 20 min, digestion was chelated with 20 mM EDTA, 4 mM EGTA, 200 mM NaCl and 1.5 pg MNase-digested *S*. *cerevisiae* mononucleosomes. Samples treated with antibodies targeting TFs, V5, or RNAPII were incubated at 4°C for 1 h to facilitate release of fragments from bead bound nuclei. The salt concentration of low salt treated samples were increased to 500 mM NaCl. Genomic fragments were then released after RNase A and Proteinase K treatment and separated through centrifugation. Fragments isolated were used as an input for library build consisting of end repair (NEBNext Ultra II End Prep Kit) and adenylation, adapter ligation, repair using USER enzyme, and subsequent purification with AMPure XP beads (Beckman Coulter). Fragments were barcoded and amplified using high fidelity polymerase (KAPA HIFI polymerase), 1.5 µM NEBNext i5 Universal PCR primers and 1.5 µM NEBNext i7 PCR primers. Samples were then purified with AMPure XP beads and eluted in 30 µL of ultrapure water. Samples were quantified by Qubit using the Qubit High Sensitivity dsDNA kit. Libraries were pooled and sequenced on an Illumina NextSeq2000 platform to a depth of ∼10million uniquely mapped reads.

### Micrococcal nuclease (MNase) digestion

MNase-seq was performed as previously described^54,90,91^. Cells were treated with either dTAG-13 or 0.1% DMSO for 2 h to deplete FACT. Cells were then counted and harvested (∼5 million cells), crosslinked with 1% formaldehyde in PBS for 10 min at room temperature in a volume of 8 mL. Cross-linking was quenched by adding 2 mL of 2.5 M Glycine, mixed thoroughly by inversion and left at room temperature for 5 min. Cells were then pelleted and rinsed twice with PBS, resuspended in cell lysis buffer (10 mM Tris pH 7.5, 10 mM NaCl, 2 mM MgCl2, 0.5% NP-40, freshly added EDTA protease inhibitors and 3 µM CaCl_2_). Micrococcal nuclease (MNase) enzyme (Takara) was aliquoted into three pre-chilled tubes in amounts of 0U, 60U and 60U. Lysed cells were added in equal amounts to each prepared tube and chromatin was digested at 37°C for 5 min. The MNase digestion was chelated with 10 mM EDTA, and samples were treated with RNAase A for 2 h at 37°C before adding 0.01% SDS and Proteinase K and incubating overnight at 55°C to reverse crosslinks. Samples were then purified using a phenol-chloroform-isoamyl alcohol (PCI) extraction, followed by extraction with chloroform and ethanol precipitation. Samples were resuspended in nuclease free-water, quantified using the Qubit High Sensitivity dsDNA kit. For traditional MNase, a small amount of the sample was run on a 1.5% agarose gel to verify sample quality. To enrich for OLDNs, the complete amount of one of the 60U digested sample was run on a 1.5% agarose gel to extract the section of the gel corresponding to 200-300 bp fragments. Paired end libraries of the MNase-digested DNA were prepared as described in the CUT&RUN section. Libraries were sequenced on a NextSeq2000 to a depth of ∼60 million reads per sample.

### MNase-ChIP

MNase ChIP was performed as described previously^54^. Briefly, cells were treated with dTAG-13 to deplete FACT or 0.1% DMSO for 2 h. Cells (∼ 5 million) were harvested and cross linked using 1% formaldehyde for 15 min at room temperature. The reaction was then quenched with 2.5 M glycine, pelleted and washed twice with PBS. Samples were then lysed with MNase (20U) for 5 min and quenched with 0.5 M EDTA. Samples were pre-cleared with protein A beads (NEB) in complete immunoprecipation buffer (20 mM Tris-HCl pH 7.5M, 1 M NaCl, 2 M EDTA, 0.1% Triton X-100 and freshly added protease inhibitor).

Input samples were taken from pre-cleared chromatin. Pre-coupled antibody-protein A beads were added to chromatin and incubated overnight at 4°C with rotation. Antibodies used were (OCT4; Invitrogen 701756 at 1:10 and SOX2; Active Motif 39843 at 1:10).Beads bound with sample were washed twice with low salt wash (20 mM Tris HCl pH 8.0, 2 mM EDTA 150 mM NaCl, 1% Triton X-100 and 0.1% SDS) and then washed twice in high salt wash (same as low salt except for NaCl which was increased to 500 mM). Samples were eluted at 65°C for 1.5 h in elution buffer (100 mM NaHCO3 and 1%SDS) and cleaned using AMPure XP beads. Paired end libraries of the DNA were prepared as described in the CUT&RUN section. Libraries were sequenced on a NextSeq2000 to a depth of ∼30 million uniquely mapped reads per sample.

### Transient transcriptome sequencing

TT-seq was performed with some modifications from a previously described method^83,92^. Cells were depleted of the FACT complex subunits for 10 min, 30 min or 2 h using dTAG-13. Following protein depletion, cells were treated with 2i LIF media containing 500 µM 4-thiouridine (4sU; Carbosynth T4509) dissolved in 100% DMSO and incubated at 37°C and 5% CO_2_ for 5 min to label nascent transcripts. After washing cells with 1X PBS, cells were pelleted and RNA was extracted using TRIzol, purified by chloroform-isopropanol extraction and precipitated with ethanol. RNA concentration was determined by Qubit RNA broad range quantification kit. Total RNA (100 µg) was diluted to a concentration of 240 ng/µl at a volume of 416.67 µl in 1X TE and fragmented with a Bioruptor Pico (Diagenode) on high power for one 30 second cycle. The fragmented RNA was combined with 10X Biotinylation Buffer (10 mM Tris pH 7.4 and 10 mM EDTA) and 200 µl of 1 mg/mL freshly prepared biotin-HPDP dissolved in dimethylformamide. Samples were incubated in a thermomixer at 37°C shaking at 1000 RPM in the dark for 2 hours to carry out Thiol-specific biotinylation. RNA was then chloroform extracted, isopropanol/ salt precipitated and dissolved in 22 µL of nuclease free water. Labeled RNA was separated from unlabeled RNA using a streptavidin C1-bead based pulldown. In brief, beads were washed in bulk in 1 mL of 0.1 NaOH + 50 mM NaCl and resuspended in binding buffer (10 mM Tris-Cl pH 7.4, 0.3 M NaCl, 1% Triton X-100) and bound to RNA for 20 min on the rotator at room temperature. Beads bound labelled RNA were washed twice with high salt buffer (5 mM Tris-Cl pH 7.4, 2 M NaCl, 1% Triton X-100), twice with binding buffer and once in low salt buffer (same as high salt but without NaCl). Nascent RNA was recovered from beads using two elutions with freshly diluted 100mM dithiothreitol at 65°C for 5 min with 1000 RPM shaking. Recovered nascent RNA was extracted with PCI and chloroform, then precipitated with ethanol. Labelled RNA was quantified using the Qubit RNA high sensitivity kit.

Strand specific nascent RNA-seq libraries were built using the NEBNext Ultra II Directional library kit (NEB) with some modifications. 100 ng of fragmented labelled RNA was used as an input for rRNA depletion via 0.05 µM pooled antisense tiling oligonucleotides followed by digestion with thermostable RNase H (MClabs). Samples depleted of rRNAs were treated with Turbo DNase (ThermoFisher) and purified by a silica column (Zymo RNA Clean and Concentrator). RNA was then fragmented at 95°C for 15 min. Fragmented RNA was used as input for cDNA synthesis and strand specific library building. NEBNext Adaptors and primers were diluted 1:2.5 and 7 PCR amplification cycles were performed for all TT-seq library builds. Libraries were quantified using Qubit with the dsDNA High Sensitivity kit and run on a fragment analyzer (Agilent) to confirm high quality of each library before sequencing. Libraries were pooled and sequenced on Illumina NextSeq2000 to a depth of ∼30 million uniquely mapped reads per sample.

### Fiber-seq^4950^

Cells were depleted of SPT16 or SSRP1 with dTAG-13 or 0.1% DMSO for 2 h. After treatment, cells were washed with 1X PBS, trypisinzed and counted at ∼ 2 million cells per aliquot. Nuclei extraction was performed by resuspending cells in 120 μL Buffer A (Freshly prepared and stored at 4°C, 15 mM Tris-HCl pH 8.0, 15 mM NaCl, 60 mM KCl, 1 mM EDTA pH 8.0 and 0.5 mM Spermidine). Samples were gently mixed by pipetting and incubated on ice for 10 min. Nuclei were pelleted (600 g, 3 min at 4°C), supernatant was discarded and nuclei was resuspended in 75 μL of Buffer A. Nuclei were then counted using a cell counter. Based on nuclei count, 1 million nuclei were transferred to a PCR tube and volume was brought up to 113 μL with Buffer A. The methyltransferase reaction was initiated by adding 3 μl of 32 mM S-adenosyl-L-methionine (SAM; final concentration of 0.8 mM) and 4 μL of Hia5 methyltransferase (EpiCypher, 15-1032) resulting in a final volume of 120 μL. Samples were mixed by gentle pipetting and incubated in a thermocycler at 25°C for 10 min. The reaction was terminated by addition of 6 μL of 10% SDS (final concentration 1%), followed by vortexing to mix. Genomic DNA was purified using Monarch Spin gDNA Purification Kit (NEB, T3010) according to the manufacturer’s instructions with minor modifications. Briefly, 400 μL of gDNA Binding Buffer was added to each reaction and mixed thoroughly by pulse-vortexing for 5-10 sec. The lysate was transferred to a gDNA purification column and centrifuged at 1000 ×g for 3 min followed by a 1 min spin at max speed. Flow-through was discarded. Columns were washed twice with 500 μL of gDNA Wash Buffer and centrifuged for 1 min at max speed after each wash. To elute DNA, 100 μL pre-warmed (60°C) gDNA Elution Buffer was added to the column, incubated at RT for 1 min and centrifuged at max speed for 1 min. DNA concentration was determined using Qubit with the dsDNA High Sensitivity kit.

Purified DNA (85 μL) was sheared using Covaris g-TUBE devices (Covaris; #PR520079). Samples were brought to a final volume of 150 μL with gDNA Elution Buffer and transferred to the screw top tube side of the g-TUBE. Samples were centrifuged at 4200 rpm for 1 min, tubes were inverted and centrifuged again at 4200 rpm for 1 min. Sheared DNA was collected from the screw-cap reservoir and transferred to a 1.5 mL Eppendorf tube. A small quantity of the samples was run on a 1% agarose gel to confirm shearing. Libraries were prepared using PacBio SMRTbell prep kit 3.0 (PacBio, 103-306-300). First, gDNA was purified using SMRTbell cleanup beads at a 1:1 bead-to-sample ratio. Briefly, 140 μL of clean up beads were added to 140 μL sheared DNA, incubated at RT for 10 min, and separated on a magnetic rack. Bound beads were washed twice with freshly prepared 80% ethanol and residual ethanol was removed by a brief spin down and separation on the magnetic rack. Beads were resuspended in 47 μL of low-TE buffer and incubated at RT for 5 min. Samples were separated on a magnetic rack and 46 μL of eluted DNA was transferred to PCR tubes. End repair/A-tailing was performed by adding 8uL Repair buffer, 4 μL End repair mix, 2 μL DNA repair mix to each sample. Samples were mixed thoroughly by pipetting and incubated at 37°C for 30 min, 65°C for 5 min. SMRTbell adapter ligation was performed using 4 μL indexed adapter, 30 μL ligation mix, 1 μL ligation enhancer and incubated at 20°C for 30 min. Ligation reactions were purified by addition of equal volume of SMRTbell cleanup beads, incubated at RT for 10min, separated on magnetic rack, and washed twice with 80% ethanol. DNA was eluted in 40 μL elution buffer. Libraries were treated with 5 μL nuclease buffer, 5 μL nuclease mix and incubated at 37°C for 15 min. Libraries were cleaned up with AMPure XP beads (0.65X) and eluted in 26 μL elution buffer. Libraries were subjected to single-molecule sequencing on PacBio Revio. All samples yielded coverage between 20-40X and an average fiber length of ∼10kb

## QUANTIFICATION AND STATISTICAL ANALYSIS

### CUT&RUN data analysis

Paired-end fastq files were trimmed to 25bp and mapped to the mm10 genome using bowtie2^93^ (options -q -N 1 -X 1000). Mapped reads were filtered from duplicates using Picard^94^ and for mapping quality (MAPQ ≥□10) using SAMTools^95^. Size classes corresponding to each factor footprints (1-120 for TFs, 1-200 for FACT, 1-120 for RNAPII, 150-500 for histone modifications) were generated using SAMTools^95^. Reads were converted to bigWig files using deepTools^96^ with the TPM-related read normalisation RPGC (options --bs 5 normalizeUsing RPGC, --effectiveGenomeSize 2308125349). For spike in normalization, scaling factor was calculated as previously described (10,000000/ uniquely mapped *Drosophila* reads) for each sample and bigwig files were generated using deepTools^96^ (option –scaleFactor). Heatmaps and metaplots were generated using deepTools^96^ computeMatrix, plotHeatmap and plotProfile. Browser track images were generated using the Integrative Genomics Viewer (IGV)^97^. SOX2 (using –style factor) and H3K4me3 (using --style histone) motifs were called and annotated using HOMER^98^. Genome-wide SOX2 motifs were called using HOMER’s command scanMotifGenomeWide.pl with SOX2 PWM from HOMER^98^ (Sox2.motif) and outputted as a bed file. Bed files were converted to binary BigWigs using UCSC BigWig and BigBed^99^ tools indicating position information of motifs which was plotted over active genes using deepTools^96^ computeMatrix, plotHeatmap and plotProfile. All CUT&RUNs were performed with 2 biological replicates for SSRP1-dTAG depletion and 2 technical replicates for SPT16-dTAG depletions. The number of genomic locations visualised in all plots are indicated with “n” under each figure. Summed occupancy analyses were quantified as a summed AUC proxy and visualised as a violin plot. Analyses and visualisation were performed in R using tidyverse, ggplot2/ggpubr, and rstatix. Changes in SOX2 occupancy and H3K4me3 occupancy were quantified across matched genomic locations by computed log_10_ scaled differences relative to DMSO ((Δlog_10_ = log_10_(signal+1) − log_10_(DMSO+1)). Mean ΔSOX2 and ΔH3K4me3 values and their standard deviations were summarized for each time point was visualised in a scatter plot. Analysis was performed in R, using data.table, tidyverse, ggplot2/ ggpubr.

### MNase-seq data analysis

Paired-end fastq files were trimmed to 25 bp and mapped to the mm10 genome using bowtie2^93^ (options -q -N 1 -X 1000). Mapped reads were filtered from duplicates using Picard^94^ and for mapping quality (MAPQ ≥□10) using SAMTools^95^. Reads were then sorted into OLDN (230-270 bp) or mononucleosome (135-165 bp) using SAMTools^95^. Sized reads were converted to bigwig files using deepTools^96^ bamCoverage (options -bs5 –normalizeusingRPGC, --effectiveGenomeSize 2308125349). Heatmaps and metaplots were generated using deepTools computeMatrix, plotHeatmap and plotProfile. MNase-digestion was performed for 1 biological replicate of SPT16 and SSPR1-dTAG depletion.

### MNase ChIP data analysis

Paired-end fastq files were trimmed to 25 bp and mapped to the mm10 genome using bowtie2^93^ (options -q -N 1 -X 1000). Mapped reads were filtered from duplicates using Picard^94^ and for mapping quality (MAPQ ≥□10) using SAMTools^95^. Reads were converted to bigWig files using deepTools with the TPM-related read normalization RPGC (options --bs 5 normalizeUsing RPGC, --effectiveGenomeSize 2308125349). Heatmaps and metaplots were generated using deepTools^96^ computeMatrix, plotHeatmap and plotProfile.

### TT-seq data analysis

Paired end fastq files were trimmed and filtered, then aligned to the mm10 genome using STAR^100^ (options -outSAMtype SAM outFilterMismatchNoverReadLmax 0.02 – outFilterMultimapNmax 1). Feature counts were generated using subread featureCounts^101^ (options -s 2 -p -B -t exon) for genes. No filtering for baseline expression was applied due to the sensitivity of TT-seq in detecting lowly expressed transcripts. To visualize TT-seq data, bigwigs were generated using deepTools^96^ with TPM read normalization (options -bs 1 –normalizeUsing BPM). Reads were imported to R and downstream analysis was conducted using DESeq2^102^ for all available replicates (n=2 for each depletion) only keeping features with at least 10 counts in 50% of samples and with option lfcShrink type = “apeglm”(DEGs reported in Supplementary file 2). Differentially expressed transcripts were plotted using EnhancedVolcano. Significance was defined as DESeq2 adjusted p-value <0.05. Strand specific bigwigs were generated with deepTools in 5 bp bins, and averaged bigwigs were generated using WiggleTools and for all replicates for each condition.

### mRNA-seq and analysis

Cells were depleted of SSRP1 for 30 min or 2 h using dTAG-13. Following protein depletion, cells were washed with 1X PBS and pelleted. RNA was extracted using Trizol, purified by chloroform-isopropanol extraction and precipitated with ethanol. RNA concentration was determined by Qubit RNA broad range quantification kit. Total RNA was subjected to Poly-A enrichment. Purified steady-state RNA samples were used as an input for strand specific RNA-seq library builds using Novogene. Libraries were pooled and paired end sequencing was performed using Illumina Novaseq platform to a depth of ∼30 million reads.

Paired end fastq files were trimmed and filtered, then aligned to the mm10 genome using STAR^100^ (options -outSAMtype SAM outFilterMismatchNoverReadLmax 0.02 – outFilterMultimapNmax 1). Feature counts were generated using subread featureCounts (options -s 2 -p -B -t exon) for genes. To visualize RNA-seq data, bigwigs were generated using deepTools with TPM read normalization (options -bs 1 –normalizeUsing BPM). Reads were imported to R and downstream analysis was conducted using DESeq2^102^ for all available replicates (n=2 for each depletion) only keeping features with at least 10 counts in 50% of samples and with option lfcShrink type = “apeglm”. Differentially expressed transcripts were plotted using EnhancedVolcano. Significance was defined as DESeq2 adjusted p-value <0.05. Strand specific bigwigs were generated with deepTools binsize of 5, and averaged bigwigs were generated using all replicates for each condition.

### Fiber-seq analysis

PacBio Fiber-seq reads were aligned to the mm10 reference genome using pbmm2 followed by indexing with SAMTools. Downstream analysis was performed using fibertools-rs/0.8.0^103^.

Samples had PacBio kinetic tags and were processed with fibertools (ft predict-m6A) to infer m6A signal, add nucleosome, and MTase sensitive patches. Quality control was performed using ft validate command. For downstream genome-browser visualization, ft pileup was used to generate per base pileup of Fiber-seq data. BED representations of m6A, 5mC and nucleosomes were generated, converted converted to bedGraph, coordinate-sorted, and written as bigwig tracks using UCSC bedGraphToBigWig^99^. To identify accessible regulatory regions, aligned BAMs were further processed using FIRE^53^ (Fiber-seq Inferred Regulatory Elements) pipeline executed via a Snakemake workflow (pixi environment), using mm10 FASTA reference, and chromosome size files. FIRE identifies regulatory elements by defining methytransferease-sensitive patches (MSPs) on individual Fiber-seq reads and then classifying MSPs as single-molecule FIRE elements as described previously^53^. FIRE peaks were called as local maxima of the FIRE score with FDR <0.05. Wide FIRE peaks were computed as the union of FIRE peaks and all regions below the FIRE threshold, followed by merging adjacent regions separated by ≤147 bp (one nucleosome). Peaks were loaded into IGV for visualization.

### Statistics

Statistical details for each experiment shown can be found in the accompanying figure legends. All experiments were performed with 2 clones for SSPR1-dTAG depletion and 1 clone for SPT16-dTAG depletions. Statistical tests were used in RNA-seq and TT-seq as per the default parameters for DESeq2 with a correction applied to minimize fold change of lowly expressed transcripts (LFCshrink). Adjusted p-values were used for significance cut off (0.05) for differential genes. Motif analyses using HOMER^98^ and peak calling were performed using default program parameters. Any error bars shown represent standard deviation in both directions. Average values for CUT&RUN, MNase-seq, and MNase-ChIP were determined by computing the mean coverage at each base pair throughout the indicated genome area between replicates. All statistical comparisons were performed in GraphPad PRISM 9 or R using rstatix. Correlative analyses were performed to compare replicates between NGS datasets for similarities and are displayed in Supplementary file 3.

## RESOURCE AVAILIBILITY

## Lead contact

All information and resource request for resources should be directed to the lead contact, Sarah Hainer.

## Materials availability

All unique materials and resources generated from this study are available upon request from the lead contact.

## Data and code availability

- Raw data and processed files will be made available upon publication on GEO.
- Any further needed information is available upon request from the lead contact.

## Acknowledgements

We thank members of the Hainer Lab for critical comments, discussion, and feedback regarding this article. We would like to thank Karen Arndt, Craig Kaplan, and Miler Lee, along with members of their labs, for discussion regarding this research. In particular we would like to thank Aakash Grover for insight on the MLN compound. We would like to thank Tom Milne and Ana Dopico-Fernandez for discussing this work prior to publication. We would like to thank the de Wit lab for kindly sharing the SOX2-dTAG and NANOG-dTAG cell lines. We would like to thank Epicypher, especially Michael Keogh and Connor Frasier, for sharing the Hia5 enzyme and Fiber-seq protocols. This work was supported by the NIH grant R35GM133732 (to S.J. Hainer). This research was supported in part by the University of Pittsburgh Center for Research Computing, RRID:SCR_022735, through the resources provided. This work used the HTC cluster, which is supported by NIH award number S10OD028483. This project used the University of Pittsburgh HSCRF Genomics Research Core, RRID: SCR_018301 next generation sequencing services, with special thanks to the Assistant Director, Will MacDonald. Research in A.G.’s laboratory was supported by the Danish National Research Foundation (DNRF195), the Novo Nordisk Foundation (NNF21OC0067425), the European Research Council (ERC-AdG no.101142230), and Independent Research Fund Denmark (DFF3101-00149B). Research at Novo Nordisk Foundation Center for Protein Research is supported by the Novo Nordisk Foundation (NNF14CC0001, NNF24SA0098829).

## Author Contributions

R.S. and S.J.H. conceived the project and designed all experiments. V.F. and A.G. designed and constructed the SSRP1-dTAG cell line and shared the cell line. B.J. generated SPT16-dTAG ES cell line and performed rescue MLN experiments. S.M.L. performed a subset of the CUT&RUN experiments. R.S. performed all other experiments. R.S. analyzed all datasets. R.S. and S.J.H. interpreted the data and wrote the manuscript. S.J.H. provided supervision. S.J.H. provided funding and A.G. provided funding for the SSRP1-dTAG cell line generation. All authors reviewed the manuscript.

## Declaration of Interests

The authors declare no competing interests.

## Notes

### Competing Interest Statement

The authors have declared no competing interest.

