## Supplemental Materials for "FACT depletion results in a temporal cascade of chromatin disruption preceding transcriptional collapse in stem cells"

**SUPPLEMENTARY MATERIALS:**

**SUPPLEMENTARY FIGURES 1-7**

**SUPPLEMENTARY TABLE 1**

**SUPPLEMENTARY FIGURES:**

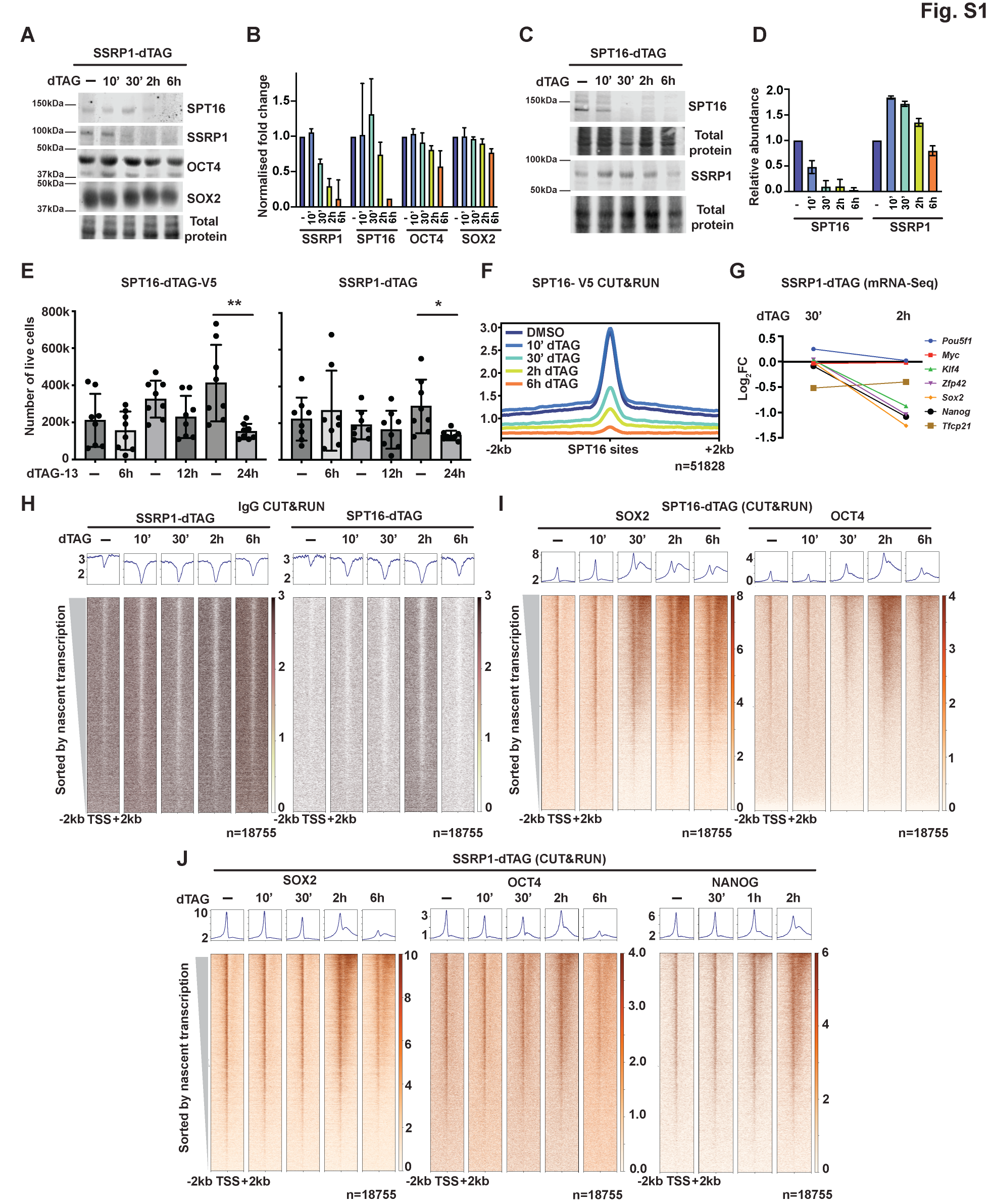

**Figure S1. Characterization of FACT depletion kinetics and effects on transcription factor localization, related to Figure 1**

**(A)** Western blot analysis of SSRP1-dTAG ES cells showing SPT16, SSRP1, OCT4, and SOX2 protein levels following dTAG-13 treatment for indicated times. Total protein (REVERT stain) serves as a loading control.

**(B)** Quantification of protein levels from (B), normalized to total protein. Data represent individual normalized fold change ± SD (n = 2 biological replicates).

**(C)** Western blot analysis of SPT16-dTAG ES cells showing SPT16 and SSRP1 protein levels following dTAG-13 treatment for indicated times. Total protein (REVERT) serves as a loading control. SPT16 depletion results in co-depletion of SSRP1.

**(D)** Quantification of protein levels from (D), normalized to total protein. Data represent mean individual normalized fold change ± SD (n = 2 biological replicates).

**(E)** Cell viability in SPT16-dTAG (left) and SSRP1-dTAG (right) ES cells following dTAG-13 treatment for indicated times. Data represent live cell counts ± SEM. *p < 0.05, **p < 0.01 (One way ANOVA, Kruskal-Wallis test for multiple comparisons).

**(F)** Metaplot of SPT16-V5 CUT&RUN enrichment at previously established FACT binding sites across the dTAG-13 treatment time course[^39^](https://sciwheel.com/work/citation?ids=15342633&pre=&suf=&sa=0&dbf=0).

**(G)** mRNA-seq showing pluripotency gene expression (*Pou5f1*, *Myc, Klf4, Zfp42, Sox2, Nanog, Tfcp21*) in SSRP1-dTAG ES cells following 30 min or 2 h dTAG-13 treatment. Data are shown as log₂ fold change relative to vehicle control (−; DMSO).

**(H)** IgG control CUT&RUN in SSRP1-dTAG (left) and SPT16-dTAG (right) ES cells. Heatmaps show signal at TSSs ± 2 kb across the dTAG-13 treatment time course. Genes (n = 18,755) are sorted by nascent transcription levels[^82^](https://sciwheel.com/work/citation?ids=17921419&pre=&suf=&sa=0&dbf=0). Metaplots (top) show average signal, confirming low background.

**(I)** Heatmaps of SOX2 (left) and OCT4 (right) CUT&RUN signal at TSSs ± 2 kb in SPT16-dTAG ES cells across the dTAG-13 treatment time course. Genes (n = 18,755) are sorted by nascent transcription levels[^82^](https://sciwheel.com/work/citation?ids=17921419&pre=&suf=&sa=0&dbf=0). Metaplots (top) show average signal.

**(J)** Heatmaps of SOX2 (left), OCT4 (middle), and NANOG (right) CUT&RUN signal at TSSs ± 2 kb in SSRP1-dTAG ES cells across the dTAG-13 treatment time course. Genes (n = 18,755) are sorted by nascent transcription levels[^82^](https://sciwheel.com/work/citation?ids=17921419&pre=&suf=&sa=0&dbf=0). Metaplots (top) show average signal.

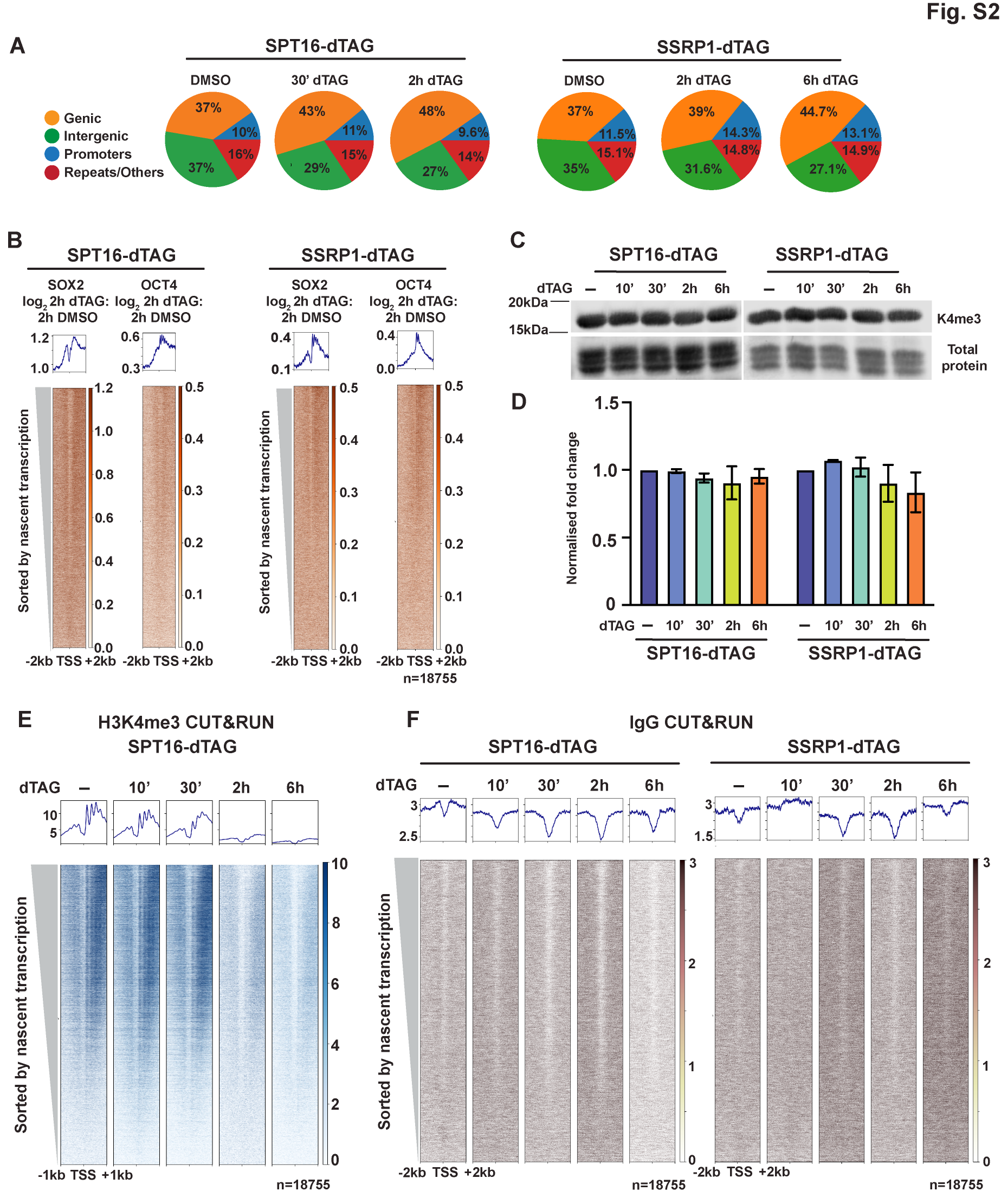

**Figure S2. FACT depletion alters TF and H3K4me3 localization, related to Figures 1 & 2**

**(A)** Pie charts showing the genomic distribution of SOX2 CUT&RUN peaks in SPT16-dTAG (left) and SSRP1-dTAG (right) ES cells following vehicle (−; DMSO) or dTAG-13 treatment for the indicated times. Peak categories include genic (orange), intergenic (green), promoters (blue), and repeats/others (red).

**(B)** MNase-ChIP validation of TF redistribution. Heatmaps showing log₂ fold change of SOX2 (left) and OCT4 (right) occupancy (2 h dTAG versus 2 h DMSO) at TSSs ± 2 kb in SPT16-dTAG (left panels) and SSRP1-dTAG (right panels) ES cells. Genes (n = 18,755) are sorted by nascent transcription levels[^82^](https://sciwheel.com/work/citation?ids=17921419&pre=&suf=&sa=0&dbf=0). Metaplots (top) show average signal across all genes.

**(C)** Western blot analysis of H3K4me3 levels in SPT16-dTAG and SSRP1-dTAG ES cells following dTAG-13 treatment for indicated times. Total protein (REVERT stain) serves as a loading control.

**(D)** Quantification of H3K4me3 protein levels from (C), normalized to total protein. Data represent mean individual normalized fold change ± SD (n = 2 biological replicates).

**(E)** Heatmap of H3K4me3 CUT&RUN signal at TSSs ± 1 kb in SPT16-dTAG ES cells across the dTAG-13 treatment time course. Genes (n = 18,755) are sorted by nascent transcription levels[^82^](https://sciwheel.com/work/citation?ids=17921419&pre=&suf=&sa=0&dbf=0). Metaplot (top) shows average signal across all genes.

**(F)** IgG control CUT&RUN in SPT16-dTAG (left) and SSRP1-dTAG (right) ES cells. Heatmaps show signal at TSS ± 2 kb across the dTAG-13 treatment time course. Genes (n = 18,755) are sorted by nascent transcription levels. Metaplots (top) show average signal, confirming low background signal across all conditions.

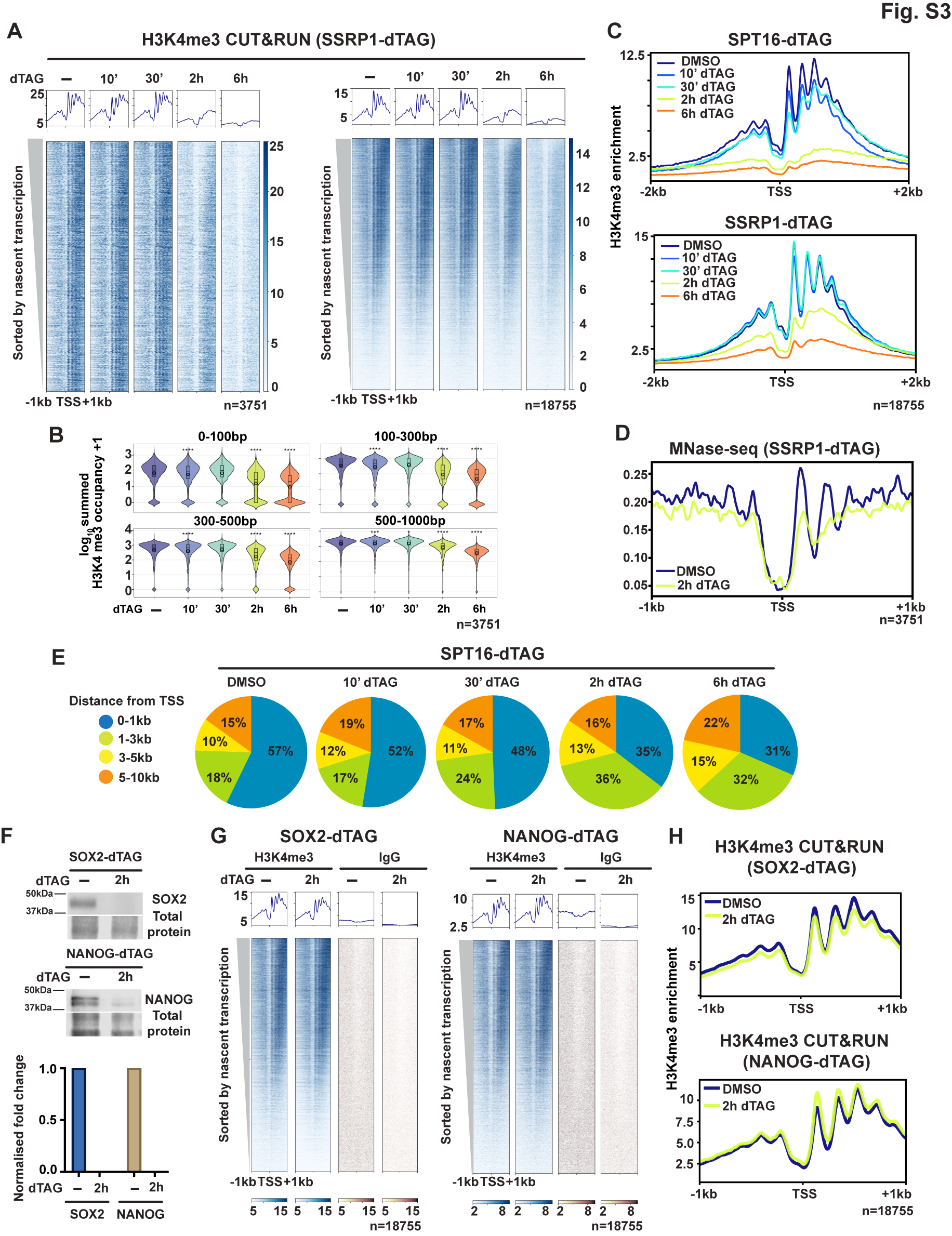

**Figure S3. H3K4me3 loss and nucleosome disruption are specific to FACT depletion, related to Figure 2**

**(A)** Heatmaps of H3K4me3 CUT&RUN signal at TSSs ± 1 kb in SSRP1-dTAG ES cells across the dTAG-13 treatment time course. Left: highly transcribed genes (n = 3,751); right: all genes (n = 18,755). Genes are sorted by nascent transcription levels[^82^](https://sciwheel.com/work/citation?ids=17921419&pre=&suf=&sa=0&dbf=0). Metaplots (top) show average signal.

**(B)** Violin plots showing summed H3K4me3 occupancy (log₁₀ + 1) across four gene body regions (0-100 bp, 100-300 bp, 300-500 bp, and 500-1000 bp downstream of TSS) in SSRP1-dTAG ES cells following dTAG-13 treatment (n = 3,751 genes). *p < 0.05, ***p < 0.001, ****p < 0.0001 (For each genomic window summed occupancy was compared across conditions using a Kruskal–Wallis test. Pairwise differences relative to DMSO were assessed using two-sided Wilcoxon rank-sum tests with Bonferroni correction; significance is indicated by asterisks).

**(C)** Metaplots of H3K4me3 CUT&RUN signal at TSSs ± 2 kb across treatment conditions in SPT16-dTAG (top) and SSRP1-dTAG (bottom) ES cells (n = 18,755 genes).

**(D)** Metaplot of MNase-seq signal at TSS ± 1 kb in SSRP1-dTAG ES cells comparing vehicle (DMSO) versus 2 h dTAG-13 treatment (n = 18,755 genes).

**(E)** Pie charts showing the genomic distribution of H3K4me3 CUT&RUN peaks with specific distances from TSSs in SPT16-dTAG ES cells following vehicle (−; DMSO) or dTAG-13 treatment for the indicated times. Peak categories include 0-1 kb from TSS (blue), 1-3 kb from TSS (green), 3-5 kb from TSS (yellow), and 5-10 kb from TSS (orange).

**(F)** Western blot analysis of SOX2 and NANOG levels in SOX2-dTAG and NANOG-dTAG ES cells following dTAG-13 treatment for indicated times. Total protein (REVERT stain) serves as a loading control (top). Quantification of SOX2 and NANOG protein levels from top, normalized to total protein. Data represent mean individual normalized fold change ± SD.

**(G)** Heatmap of H3K4me3 CUT&RUN signal at TSSs ± 1 kb in SOX2-dTAG (left) NANOG-dTAG ES cells (right) treated with vehicle (DMSO) or dTAG-13 for 2 h. IgG controls are shown in grey. Genes (n = 18,755) are sorted by nascent transcription levels[^82^](https://sciwheel.com/work/citation?ids=17921419&pre=&suf=&sa=0&dbf=0). Metaplot (top) shows average signal.

**(H)** Metaplots of H3K4me3 CUT&RUN signal at TSSs ± 1 kb in SOX2-dTAG (top) and NANOG-dTAG (bottom) ES cells comparing vehicle control (−; DMSO) versus 2 h dTAG-13 treatment (n = 18,755 genes).

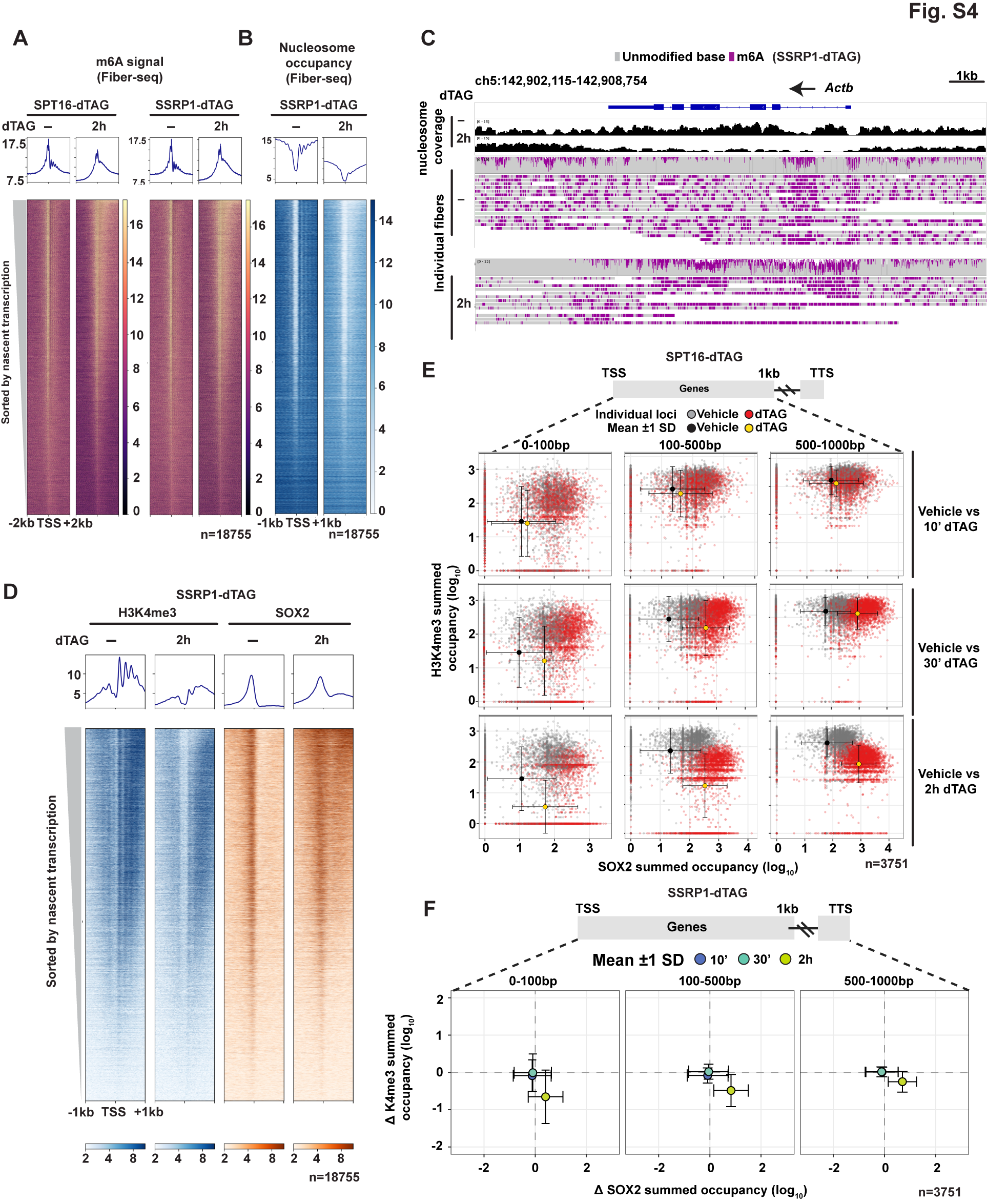

**Figure S4. H3K4me3 loss and SOX2 accumulation occur at the same genomic locations in SPT16 and SSRP1-dTAG, related to Figure 3**

**(A)** Heatmaps of m6A signal from Fiber-seq analysis at TSSs ± 2 kb in SPT16-dTAG (left) and SSRP1-dTAG (right) ES cells comparing vehicle control (−; DMSO) and 2 h dTAG-13 treatment. Genes (n = 18,755) are sorted by nascent transcription levels[^82^](https://sciwheel.com/work/citation?ids=17921419&pre=&suf=&sa=0&dbf=0). Metaplots (top) show average signal.

**(B)** Heatmap of nucleosome occupancy derived from Fiber-seq analysis at TSSs ± 1 kb in SSRP1-dTAG ES cells comparing vehicle control (−; DMSO) and 2 h dTAG-13 treatment. Genes (n = 18,755) are sorted by nascent transcription levels. Metaplot (top) shows average nucleosome occupancy.

**(C)** Chromatin fibers from Fiber-seq over the *Actb* locus (chr5:142,902,115-142,908,754). Nucleosome coverage tracks (top) and individual fiber representations are shown for vehicle control (−; DMSO) and 2 h dTAG-13 treatment. Gray indicates unmodified bases; purple indicates N6-methyladenine (m6A) marks generated by Hia5 methyltransferase, which labels open DNA.

**D)** Side-by-side heatmaps of H3K4me3 (left) and SOX2 (right) CUT&RUN signal at TSSs ± 1 kb in SSRP1-dTAG ES cells comparing vehicle control (−; DMSO) and 2 h dTAG-13 treatment. Genes (n = 18,755) are sorted by nascent transcription levels[^82^](https://sciwheel.com/work/citation?ids=17921419&pre=&suf=&sa=0&dbf=0). Metaplots (top) show average signal across all genes.

**(E)** Scatterplots comparing changes in H3K4me3 occupancy (log_10_ summed occupancy, y-axis) versus changes in SOX2 occupancy (log_10_ summed occupancy, x-axis) across three gene body regions (0-100 bp, 100-500 bp, and 500-1000 bp downstream of TSS) at 10 min (top), 30 min (middle), and 2 h (bottom) of SPT16 depletion (n = 3,751 genes). Grey and red points represent occupancy individual loci in DMSO and dTAG-13 treated conditions respectively. Black and yellow data points represent mean ± 1 SD in DMSO and dTAG-13 conditions respectively.

**(F)** Scatterplots comparing changes in H3K4me3 occupancy (Δ K4me3, relative to vehicle control, y-axis) versus changes in SOX2 occupancy (Δ SOX2, relative to vehicle control, x-axis) across three gene body regions (0-100 bp, 100-500 bp, and 500-1000 bp downstream of TSS) at 10 min (blue), 30 min (teal), and 2 h (yellow) of SSRP1 depletion (n = 3,751 genes). Data points represent mean ± 1 SD.

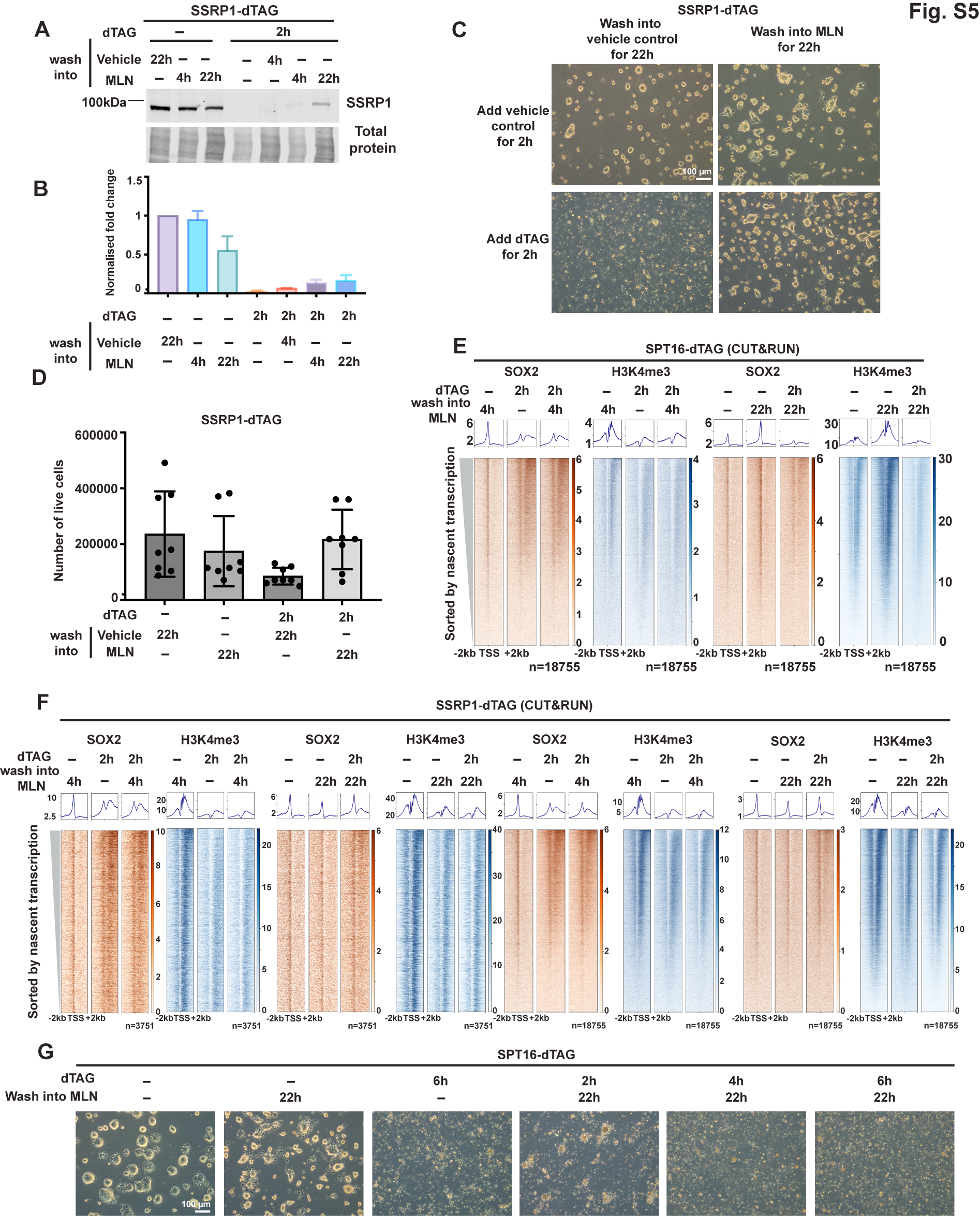

**Figure S5. FACT restoration after degradation partially rescues transcription factor mislocalization and H3K4me3 loss, related to Figure 4**

**(A)** Western blot analysis of SSRP1 protein levels in SSRP1-dTAG ES cells across treatment conditions. Cells were treated with vehicle control (−; DMSO) or dTAG-13 for 2 h, followed by washout into MLN4924 or vehicle control for 4 or 22 h. Total protein (REVERT) serves as a loading control.

**(B)** Quantification of SSRP1 protein levels from (A), normalized to total protein. Data represent normalized fold change ± SD (n = 2 biological replicates).

**(C)** Representative brightfield microscopy images of SSRP1-dTAG ES cells after 22 h of washout into vehicle control (left) or MLN4924 (right), following initial 2 h treatment with vehicle control (top) or dTAG-13 (bottom).

**(D)** Quantification of cell viability in SSRP1-dTAG ES cells across treatment conditions. (One way ANOVA, Kruskal-Wallis test for multiple comparisons).

**(E)** Heatmaps of SOX2 (left) and H3K4me3 (right) CUT&RUN signal at TSSs ± 2 kb in SPT16-dTAG ES cells across treatment conditions. Genes (n = 18,755) are sorted by nascent transcription levels[^82^](https://sciwheel.com/work/citation?ids=17921419&pre=&suf=&sa=0&dbf=0). Metaplots (top) show average signal.

**(F)** Heatmaps of SOX2 and H3K4me3 CUT&RUN signal at TSSs ± 2 kb in SSRP1-dTAG ES cells across treatment conditions at 4 h (left panels) and 22 h (right panels) washout timepoints. Highly transcribed genes (n = 3,751, left) and all genes (n = 18,755, right) are shown, sorted by nascent transcription levels[^82^](https://sciwheel.com/work/citation?ids=17921419&pre=&suf=&sa=0&dbf=0). Metaplots (top) show average signal.

**(G)** Representative brightfield microscopy images of SPT16-dTAG ES cells after 2 h vehicle washed into 22 h vehicle control, 2 h vehicle washed into 22 h MLN4924, 6 h dTAG-13 treatment washed into 18 h vehicle treatment, 2 h, 4 h and 6 h dTAG-13 treatment washed into 22 h MLN4924.

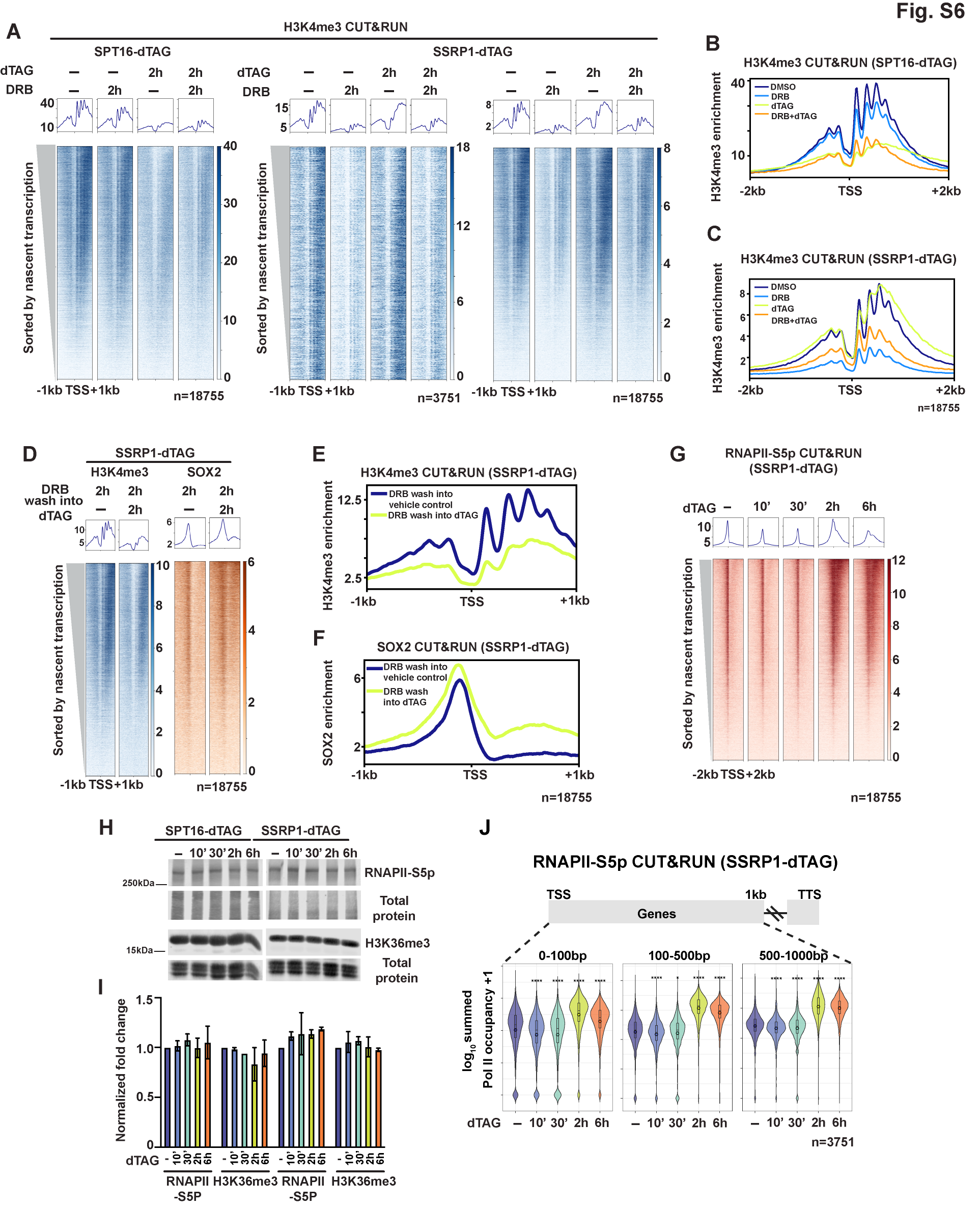

**Figure S6. Transcription-dependence of FACT-mediated chromatin maintenance and RNAPII dynamics, related to Figures 5 and 6**

**(A)** Heatmaps of H3K4me3 CUT&RUN signal at TSSs ± 1 kb in SPT16-dTAG (left, n = 18,755 genes) and SSRP1-dTAG (middle, n = 3,751 highly transcribed genes; right, n = 18,755 genes) ES cells across treatment conditions: vehicle control (−; DMSO), DRB alone, dTAG-13 alone, or DRB combined with dTAG-13 for 2 h. Genes are sorted by nascent transcription levels[^82^](https://sciwheel.com/work/citation?ids=17921419&pre=&suf=&sa=0&dbf=0). Metaplots (top) show average signal.

**(B)** Metaplot of H3K4me3 CUT&RUN enrichment at TSSs ± 2 kb in SPT16-dTAG ES cells across all four treatment conditions (n = 18,755 genes).

**(C)** Metaplot of H3K4me3 CUT&RUN enrichment at TSSs ± 2 kb in SSRP1-dTAG ES cells across all four treatment conditions (n = 18,755 genes).

**(D)** Heatmaps of H3K4me3 (left) and SOX2 (right) CUT&RUN signal at TSSs ± 1 kb in SSRP1-dTAG ES cells following DRB pre-treatment and washout into vehicle control or dTAG-13 for 2 h. Genes (n = 18,755) are sorted by nascent transcription levels[^82^](https://sciwheel.com/work/citation?ids=17921419&pre=&suf=&sa=0&dbf=0). Metaplots (top) show average signal.

**(E)** Metaplot of H3K4me3 CUT&RUN enrichment at TSSs ± 1 kb in SSRP1-dTAG ES cells comparing DRB washout into vehicle control versus DRB washout into dTAG-13 (n = 18,755 genes).

**(F)** Metaplot of SOX2 CUT&RUN enrichment at TSSs ± 1 kb in SSRP1-dTAG ES cells comparing DRB washout into vehicle control versus DRB washout into dTAG-13 (n = 18,755 genes).

**(G)** Heatmap of RNAPII-S5p CUT&RUN signal at TSS ± 2 kb in SSRP1-dTAG ES cells across the dTAG-13 treatment time course (left). Genes (n = 18,755) are sorted by nascent transcription levels[^82^](https://sciwheel.com/work/citation?ids=17921419&pre=&suf=&sa=0&dbf=0). Metaplot (top) shows average signal.

**(H)** Western blot analysis of RNAPII-S5p levels and H3K36me3 levels in SPT16-dTAG and SSRP1-dTAG ES cells following dTAG-13 treatment for indicated times. Total protein (REVERT) serves as a loading control.

**(I)** Quantification of protein levels from (H), normalized to total protein. Data represent normalized fold change ± SD (n = 2 biological replicates).

**(J)** Violin plots showing summed RNAPII-S5p occupancy (log₁₀ + 1) across three gene body regions (0-100 bp, 100-500 bp, and 500-1000 bp downstream of TSS) in SSRP1-dTAG ES cells following dTAG-13 treatment (n = 3,751 genes). *p < 0.05, **p < 0.01, ***p < 0.001, ****p < 0.0001 (For each genomic window summed occupancy was compared across conditions using a Kruskal–Wallis test. Pairwise differences relative to DMSO were assessed using two-sided Wilcoxon rank-sum tests with Bonferroni correction; significance is indicated by asterisks).

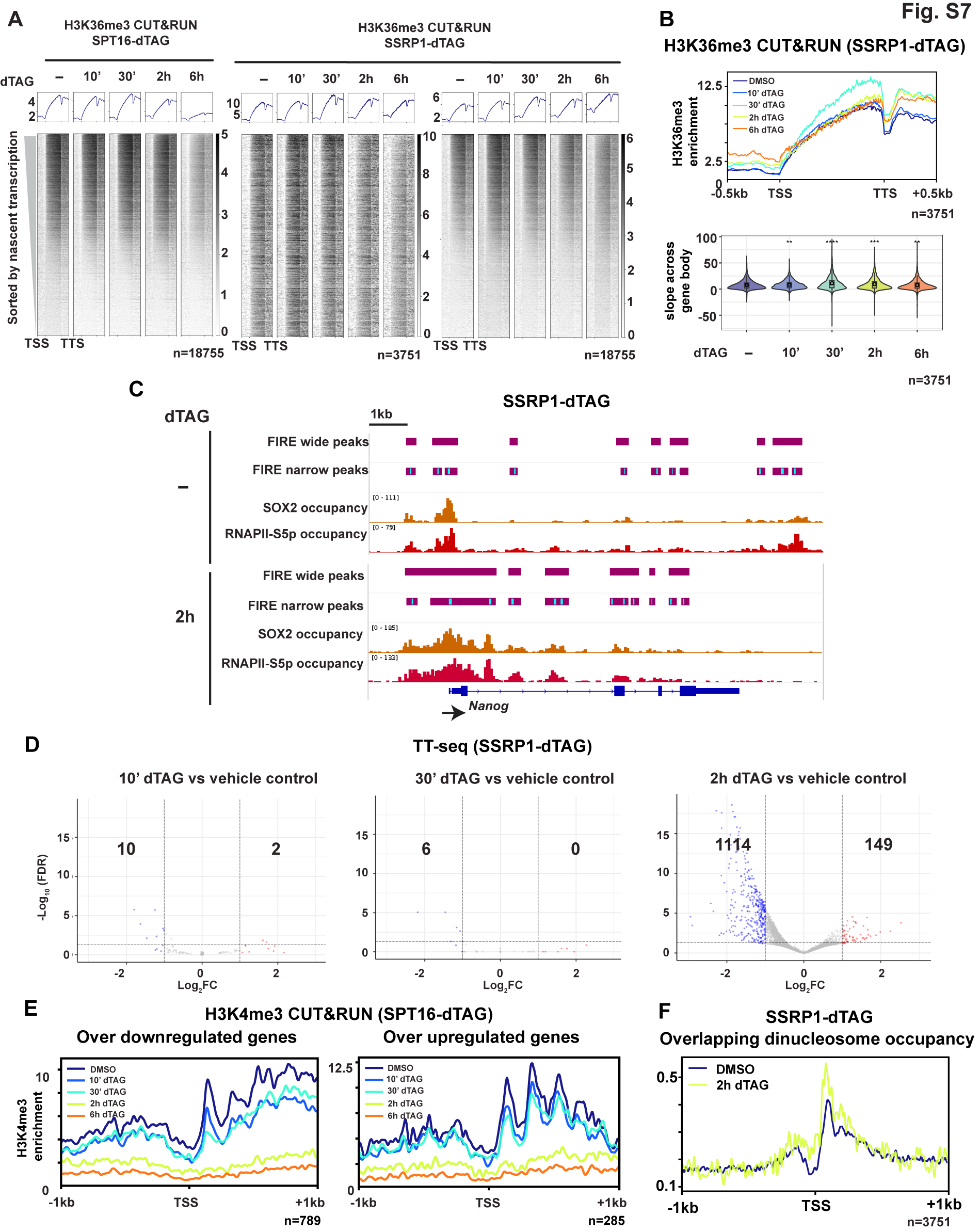

**Figure S7. RNAPII dynamics, nucleosome disruption, and transcriptional changes upon SSRP1 depletion, related to Figures 6 and 7**

**(A)** Metagene heatmaps of H3K36me3 CUT&RUN signal across gene bodies (TSS to TTS) in SPT16-dTAG (n = 18,755 genes) and SSRP1-dTAG (n = 3,751 highly transcribed genes; n = 18,755 all genes) ES cells across the dTAG-13 treatment time course (right panels). Genes are sorted by nascent transcription levels[^82^](https://sciwheel.com/work/citation?ids=17921419&pre=&suf=&sa=0&dbf=0). Metaplots (top) show average signal.

**(B)** Top: Metaplot of H3K36me3 CUT&RUN enrichment across scaled gene bodies (TSS to TTS ± 0.5 kb) for highly transcribed genes (n = 3,751) in SSRP1-dTAG ES cells across treatment conditions. Bottom: Violin plots quantifying the slope of H3K36me3 signal across gene bodies (n = 3,751 genes) following SSRP1 depletion. **p < 0.01, ***p < 0.001 (Pairwise, non-parametric Wilcoxon rank-sum, after Bonferroni corrections).

**(C)** FIRE peaks over the *Nanog* locus in SSRP1-dTAG ES cells treated with vehicle (DMSO, top) or dTAG-13 for 2 h (bottom). Tracks show (from top to bottom): FIRE-called narrow and wide peaks, FIRE-called wide peaks (nucleosome-depleted regions), FIRE score per peak, SOX2 CUT&RUN, RNAPII-S5p CUT&RUN, and gene annotation.

**(D)** Volcano plots showing differential nascent transcription at 10 min of SSRP1 depletion compared to vehicle control in SSRP1-dTAG ES cells. X-axis shows log₂ fold change; y-axis shows −log₁₀FDR. Significantly (FDR<0.05 and Log_2_FC>1) upregulated genes (red) and downregulated genes (blue) are highlighted.

**(E)** Metaplot of H3K4me3 CUT&RUN occupancy in SPT16-dTAG ES cells at TSSs ± 1 kb of significantly downregulated genes (FDR<0.05, Log_2_FC <1, n= 789, right) and upregulated genes (FDR<0.05, Log_2_FC >1, n= 285 left).

**(F)** Metaplot of overlapping dinucleosome (OLDN, 230-270 bp size class) occupancy at TSS ± 1 kb in SSRP1-dTAG ES cells comparing vehicle control (DMSO) versus 2 h dTAG-13 treatment.

**SUPPLEMENTARY TABLE 1. Table of oligonucleotide and homology constructs used in this study.**

| sgRNA for SPT16-dTAG F | CACCGAGAATGGAGCATAGGACCCA |
| --- | --- |
| sgRNA for SPT16-dTAG R | AAACTGGGTCCTATGCTCCATTCTC |
| SPT16-dTAG screening primer F | GAAGGTGCAGAGCAGTTGAGC |
| SPT16-dTAG screening primer R | AGCTTGGTCCGCACAAATGG |
| sgRNA for SSRP1-dTAG | CCTCGGGATCTGATGAA |
| SSRP1-dTAG homology F | CAGATTGTGTTTGCCAGGTCAG |
| SSRP1-dTAG homology R | GTCCGGCTGGAAAGGACAC |
| SSRP1-dTAG screening primer | TTGGCTAATGGAGTGGAGTGAAC |
| FKBP-F36V sequence used in SPT16 and SSRP1-dTAG | TTCCAGTTTTAGAAGCTCCACATCGAAGACGAGAGTGGCATGTGGTGGGATGATGCCTGGGTGCCCAGTGGCACCATAGGCATAATCTGGAGATATAGTCAGTTTGGCTCTCTGACCCACACTCATCTGGGCAACCCCTTCTTCCCAGCCTCGGATCACCTCCTGCTTGCCTAGCATAAACTTAAAGGGCTTGTTTCTGTCCCGGGAGGAATCAACTTTCTTTCCATCTTCAAGCATCCCGGTGTAGTGCACCACGCAGGTCTGGCCGCGCTTGGGGAAGGTGCGCCCGTCTCCTGGGGAGATGGTTTCCACCTGCACTCCCAT |
| V5 tag sequence | GGTAGAGTCCAGGCCGAGCAGAGGGTTAGGGATAGGTTTTCC |
